# Mechanical control of morphogenetic robustness in an inherently challenging environment

**DOI:** 10.1101/2020.01.06.896266

**Authors:** Emmanuel Martin, Sophie Theis, Guillaume Gay, Bruno Monier, Christian Rouvière, Magali Suzanne

**Affiliations:** LBCMCP, Centre de Biologie Intégrative (CBI), Université de Toulouse, CNRS, UPS, France; Morphogénie Logiciels, 32110 St Martin d’Armagnac, France; Image Processing Facility, Centre de Biologie Intégrative (CBI), Université de Toulouse, CNRS, UPS, France

## Abstract

Epithelial sheets undergo highly reproducible remodeling to shape organs. This stereotyped morphogenesis depends on a well-defined sequence of events leading to the regionalized expression of developmental patterning genes that finally triggers downstream mechanical forces to drive tissue remodeling at a pre-defined position. However, how tissue mechanics controls morphogenetic robustness when challenged by intrinsic perturbations in close proximity has never been addressed.

Here, we show that a bias in force propagation ensures stereotyped morphogenesis despite the presence of mechanical noise in the environment. We found that knockdown of the Arp2/3 complex member Arpc5 specifically affects fold directionality without altering neither the developmental nor the force generation patterns. By combining *in silico* modeling, biophysical and ad hoc genetic tools, our data reveal that junctional Myosin II planar polarity favors long-range force channeling and ensures folding robustness, avoiding force scattering and thus isolating the fold domain from surrounding mechanical perturbations.

## Introduction

An intriguing characteristic of biology is the remarkable reproducibility in size and shape of a given structure from one individual to another. This trait, called robustness, is characterized by a low level of variation of a given phenotype when subjected to environmental or genetic variations (Waddington, 1942).

An important factor ensuring robustness is the existence of redundant mechanisms that appeared to be frequently used as a way to ensure the generation of a specific trait. It has been observed at different levels, between different genes, but also between different regulatory sequences governing gene expression with the discovery of shadow enhancers or more recently between mechanical networks governing tissue shape generation (Frankel et al., 2010; Perry et al., 2010; Smith et al., 2018; Yevick et al., 2019). These discoveries highlight the importance of backup mechanisms to ensure a correct shape. However, they did not inform on the process of canalization of a particular trait and the protection mechanisms taking place to avoid phenotypic variation in front of environmental challenges.

The establishment of precise patterning constitutes the initial step for the development of anatomical structures. Not surprisingly, it has been the main focus in the field to unravel the mechanisms responsible for the high level of precision observed in terms of growth, scale and patterning of tissue and organs (Félix and Barkoulas, 2015; Lander, 2011). Pioneer studies came from the study of signaling gradients in Drosophila, highlighting the high level of precision of morphogen gradients giving rise to precise boundary of target genes expression, even when challenged by fluctuation in gene dosage or morphogen production rate (Eldar et al., 2004; Gregor et al., 2007; Hardway et al., 2008; He et al., 2008). It then became apparent that even if morphogen gradients appear amazingly robust, a certain degree of variability exists from one cell to another. This stochasticity in molecular and cellular processes emerged recently as an inherent part of biological systems, opening a whole field of research focused on how a developing organism can deal with such intrinsic noise and form stereotyped shapes (Ebert and Sharp, 2012; Gursky et al., 2012; Hong et al., 2018; Lander, 2011).

Numerous studies highlighted the importance to buffer this noise. Indeed, even in situations where, counter intuitively, heterogeneity has been shown to play an important role in shape robustness such as in sepal formation, heterogeneity has to be buffered over time to ensure the formation of regular and stereotyped organs (Hong et al., 2018). Different mechanisms of noise buffering have been identified, mainly involved in fine-tuning the expression of genes involved in positional information (Ebert and Sharp, 2012; Gursky et al., 2012; Herranz and Cohen, 2010; Lott et al., 2007; Manu et al., 2009; Sato, 2018).

Downstream of this positional information and the establishment of tissue coordinates, important drivers of tissue shape are mechanical forces, which govern cell shape remodeling and cell rearrangement processes (Smith et al., 2018). The study of mechanical forces constitutes a growing field in morphogenesis, underlying their crucial role in tissue dynamics. Indeed, cells and tissues are physical entities, whose shape is determined by structural components, such as cytoskeletal proteins. Among them, the molecular motor non-muscle myosin II and the filamentous actin associate to create a dynamic network, which drives force generation or governs cell architecture. The re-distribution of acto-myosin within specific subcellular domains drives specific changes in cell shape. These forces, generated at a single cell level can propagate from one cell to another through adherens junction or diffusible biomechanical signals (Lecuit and Yap, 2015), ultimately leading to large-scale changes and tissue shape modification, although long-range propagation and channeling of mechanical forces remains largely unexplored.

If the cellular machinery responsible for force generation is a key player in the construction of a particular shape, only a few works addressed the question of mechanical contribution to robustness. On the one hand, Hong and colleagues (Hong et al., 2018) proposed, based on theoretical modeling, that tissue mechanics could buffer the heterogeneity observed at a single cell level in term of growth and stiffness. More recently, this idea has been tested experimentally and mechanical forces have been shown to buffer local heterogeneity both in zebrafish and in Drosophila (Akieda et al., 2019; Eritano et al., 2020). On the other hand, Yevick and colleagues (Yevick et al., 2019) identified redundant mechanical networks involved in the construction of a particular shape. However, despite an important amount of works on mechanical forces as essential bricks in the construction of tissue shape, how mechanics affects the degree of variability of a given phenotype in challenging environmental conditions remain mostly unexplored. Indeed, while the generation and the propagation of forces are well recognized as an important property of cells and tissues, they are mainly viewed as key executioners of a pre-established developmental program or as a refining mechanism of intrinsic noise rather than a guardian of the stereotypic nature and robustness of tissue shape in front of external challenges.

Here, focusing on tissue invagination as a model of morphogenetic robustness, we identified a genetic condition in which fold orientation, an extremely robust trait, becomes highly variable. This variability appears to be independent of tissue patterning; with both tissue regionalization and intrinsic mechanical signals occurring normally, while fold directionality is perturbed. These deviations point at regions of high tension, which appear randomly around the fold domain, thus revealing the presence of mechanical noise in the local environment. Finally, we found that Myosin II planar polarity is both necessary and sufficient to ensure the robustness of fold directionality, favoring force channeling, and thus protecting the invagination from neighboring mechanical noise.

## Results

### Morphogenesis variability in Arpc5 knockdown

To address the role of tissue mechanics in morphogenetic robustness, we used Drosophila leg development, a model particularly appropriate since it undergoes a highly stereotyped morphogenesis with four parallel folds formed during development, in the most distal part of the leg or tarsal region (Fig1a). To quantify the robustness of this morphogenetic process in a control situation, we measured the variability of fold positioning (distance from the predicted fold), fold orientation (angle formed with the proximal-distal axis of the leg) and further quantified the relative orientation of tarsal folds between them or fold parallelism (Fig1c-f, control). Both fold positioning and orientation appear to be extremely robust in the control situation, as shown by the low standard deviation observed for each of these parameters.

**Figure 1:**
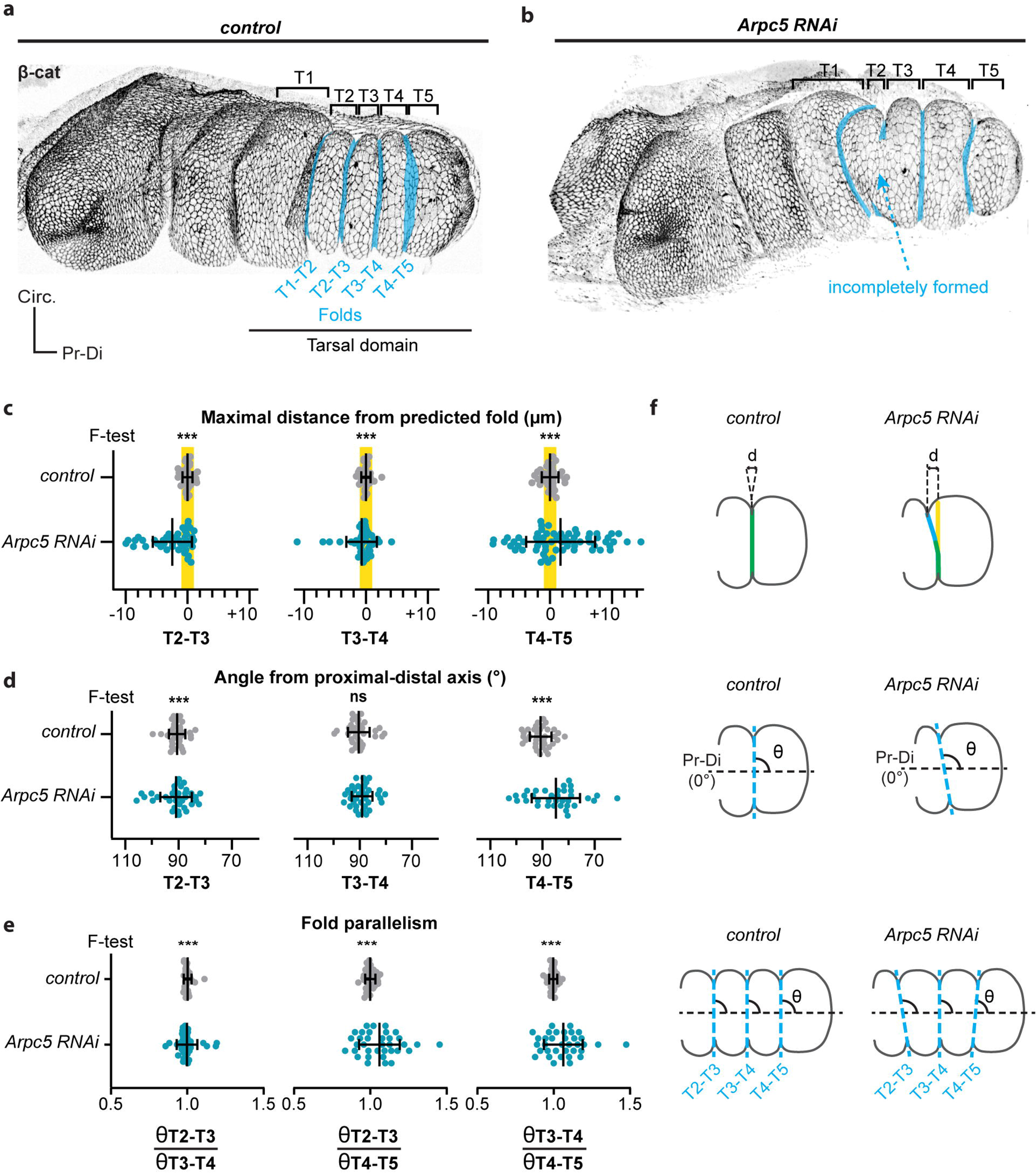
Fold morphogenetic robustness is lost in Arpc5 knockdown (related to FigS1). **a-b**, 3D reconstructions of a *control* (Dll-Gal4; arm-GFP) (a) or *Arpc5* RNAi (*Dll-Gal4; arm-GFP, UAS Arpc5RNAi*) **(b)** pupal leg discs at 1h30 after puparium formation (APF), showing tarsal folds morphology in blue. **c,** Dot plots showing the maximal distance between the real folds and the corresponding predicted fold (highlighted in yellow) in control and *Arpc5* RNAi leg discs (n=35, 63, respectively). **d,** Dot plots showing the angle of the folds relative to the proximal-distal axis in control and Arpc5 RNAi discs (n=35 in both cases). **e,** Dot plots quantifying the fold parallelism (ratio between the angles of two different folds) in *control* and *Arpc5* RNAi leg discs (n=35 in both cases). In **c-e,** a F-test of equality of variances was used. ns, not significant; ***, p-value < 0.001. Black lines represent the mean ± SD. Genotypes correspond to *sqh-RFPt[3B]; Dll-Gal4, UAS-GC3Ai* uncrossed or crossed to *UAS Arpc5RNAi*.

To decipher how this particular shape is established and maintained from one individual to another, we selected, from an unbiased screen expressing a library of RNAi in the tarsal domain of the leg (manuscript in preparation), candidates affecting specifically fold positioning. Interestingly, we found a puzzling phenotype, never described before, of misoriented folds. This phenotype was found for several components of the Arp2/3 complex known to regulate the branched actin network, including Arpc5, Arpc3a and Arp2 (Fig1b, S1b) (Chesarone and Goode, 2009; Pollard and Beltzner, 2002; Robinson et al., 2001). Interestingly, defects show a wide range of variability and ranged from a complete absence of folds to fold deviation to normal folding (FigS1b-c). Since these proteins are all part of the core Arp2/3 complex and their RNAi give similar phenotypes (FigS1b-c), we focused our analysis on one of them, Arpc5, whose inactivation by RNAi in our experimental conditions resulted in a 50% reduction of its mRNA level (FigS1a). The implication of Arp2/3 in fold directionality was unexpected. To further characterize this new phenotype, we first compared the defects observed during fold formation and at the end of the process. We observed an increase in the proportion of fold deviation over time, going from 1/3 to 3/4. These results indicate that the deviations are not transient but stable defects. We further observed that the absence of folding becomes less frequent with time, suggesting together with the increase of fold deviations that a proportion of folds are delayed and finally misoriented (FigS1c). To further characterize this mutant phenotype, we choose to focus on mid-fold formation, a stage at which fold positioning can be defined more accurately. We then measured the variability of fold positioning, fold orientation and fold parallelism of the subpopulation of discs (about 2/3) displaying normal or misoriented folds (Fig1c-f), and excluding unfolded legs since fold direction is impossible to address in these ones (mentioned as flat in FigS1c). For each of these parameters, standard deviation was significantly higher than in the control, highlighting the high degree of variability observed in the Arpc5 knockdown condition (Fig1c-f), a phenotype characteristic of a failure of morphogenetic robustness.

### Developmental patterning in Arpc5 knockdown

Although the regulation of fold orientation has never been addressed, fold positioning is known to be determined by the sequential establishment of positional information along the developing leg tissue, starting with the restricted expression of morphogens such as *wingless* and *decapentaplegic* (Lecuit and Cohen, 1997), the subsequent proximal-distal regionalization through the expression of patterning genes such as *Hth*, *Dac* and *Dll* (Wu and Cohen, 1999), followed by the segmental activation of the Notch pathway in the distal part of each segment (de Celis et al., 1998; Rauskolb and Irvine, 1999). Finally, in the tarsal region, pro-apoptotic genes are expressed in a few rows of cells in the distal part of each segment (Manjón et al., 2007). This positional information is then translated into a “patchy” pattern of apoptotic cells in the predicted fold domain. Previous work that reconstituted the localization of apoptotic cells in fixed samples over time showed that apoptotic cells appear first on the ventral part, then in the lateral domains, and finally in the most dorsal domain of this ring-shaped predicted fold (Monier et al., 2015). Before their elimination, these apoptotic cells exert apico-basal forces, which constitute the initiator mechanical signals for tissue remodeling. These forces are transmitted to their neighbors leading to an increase in apical myosin accumulation, apical constriction and eventually tissue folding, a process that lasts 3-4h *in vivo* (Monier et al., 2015).

Since robustness has been shown to rely on the establishment of robust positional information, we first analyzed the expression pattern of genes known to be involved in fold positioning. Importantly, in the Arpc5 knockdown condition, the segmental activation of Notch, characterized by the expression of the Notch target gene *Deadpan* (*Dpn*), is maintained (Fig2a). Indeed, on top of *Dpn* expression in neurons (see asterisks in Fig2a), consistent with its identification as a pan-neural gene (Younger-Shepherd et al., 1992), *Dpn* is expressed in stripes in the distal leg, both in the control and Arpc5 RNAi condition. We further characterized *Dpn* expression domain and found that the width of its domain is slightly smaller in Arpc5 RNAi (FigS2d), consistent with the role played by Arp2/3 in Notch activation in other contexts (Rajan et al., 2009). However, the orientation of *Dpn* stripes in relation to the proximal-distal axis, their parallelism and their curvature were unaffected in Arpc5 knockdown condition (Fig2c; FigS2a,c). Fold deviation in the Arpc5 knockdown appears to result from a partial dissociation of the folding process from the positional information. Thus, while folds follow the segmental stripes of Notch activation domain in the control, folds deviate from these positional cues in the Arpc5 knockdown (Fig2e-f, MovieS1 and S2). We next analyzed the apoptotic pattern in the Arpc5 knockdown, which appears intact as shown by the unperturbed expression of the pro-apoptotic gene *reaper* (Fig2b,d; FigS2b,e,f) and the frequency of apoptotic cells in the fold domain (Fig2g). Finally, the ability of dying cells to generate mechanical signals is identical to the control, as shown by the formation of apico-basal transient structures of Myosin II in dying cells (Fig2h), their ability to deform the apical surface (Fig2h) and to generate apico-basal tension (Fig2i, MovieS3). Thus, while folds deviate in Arpc5 knockdown condition, positional information resulting from the developmental patterning and the subsequent mechanisms known to be involved in the fold formation are unaffected. This surprising result prompted us to revisit the prevailing model of morphogenetic robustness relying on strict regulation of morphogen gradients or key identity genes (Gilmour et al., 2017). We hypothesized that fold deviation could either be due to the appearance of a new source of perturbation or by an increased sensitivity to existing perturbations in the knockdown condition.

**Figure 2:**
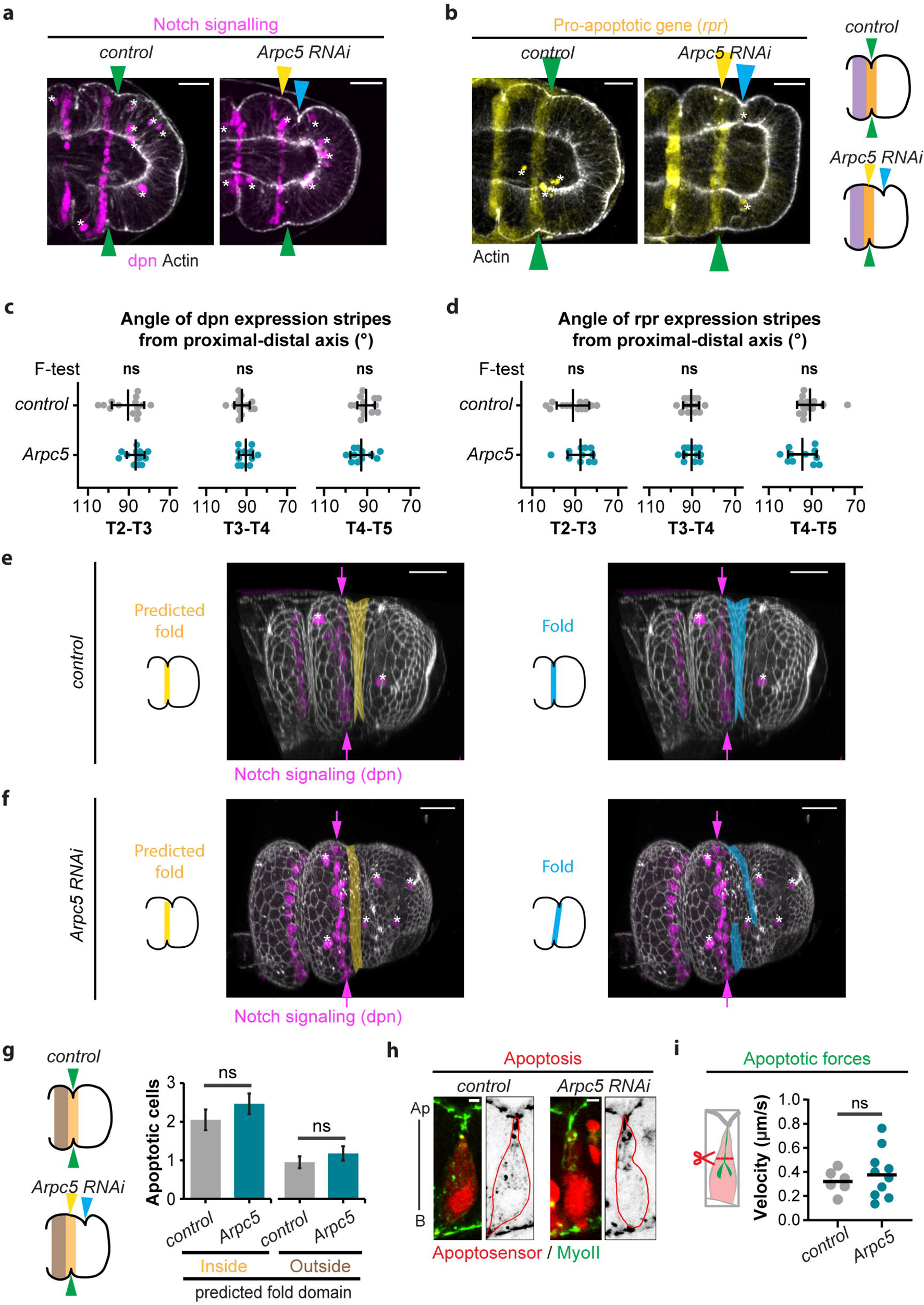
Developmental patterning is unaffected in Arpc5 knockdown (Related to FigS2 and Movie S1-S3) **a,** Z-sections of *control (sqh-GFP[29B]; Dll-Gal4)* and *Arpc5 RNAi (sqh-GFP[29B]; Dll-Gal4; UAS-Arpc5RNAi)* leg discs stained with phalloidin (white) and showing Z-projection of anti-Deadpan (magenta). **b,** Z sections of *control (rpr-lacZ; Dll-Gal4) or Arpc5 RNAi (rpr-lacZ; Dll-Gal4; UAS-Arpc5RNAi)* leg discs stained with phalloidin (white) and showing a Z-projection of rpr-lacZ (yellow). **a,b**, Yellow, blue and green arrowheads respectively indicate predicted fold, real fold and perfect match between them. Please note that both Dpn and rpr-lacZ are also expressed in some neurons, indicated by asterisks. Cartoons on the right recapitulate the location of Notch signaling (purple) and pro-apoptotic genes expression (yellow) in *control and Arpc5* RNAi conditions. **c-d**, Dot plots showing the angle of the stripes of expression of *Deadpan* (c) or *reaper* (d) in the folds relative to the proximal-distal axis in *control (rpr-lacZ; Dll-Gal4)* and *Arpc5* RNAi *(rpr-lacZ; Dll-Gal4; UAS-Arpc5RNAi)* leg discs (n=13 and 11 respectively). A *F*-test of equality of variances was used. ns, not significant. Black lines represent the mean ± SD. **e-f**, 3D-reconstructions of control (e) and *Arpc5* RNAi (f) leg discs showing the expression of Deadpan (magenta, arrows). Yellow and blue domain respectively highlight the predicted and the real fold. Asterisks point out *Dpn-expressing* neurons. **g**, Average number of dying cells inside or outside the predicted distal-most tarsal fold in control and Arpc5 RNAi leg discs (n=20 and 28, respectively). Bar graphs indicate the mean ± SEM. Cartoons on the left recapitulate the regions inside (yellow) and outside (brown) of the predicted fold. **h,i**, Sagittal views showing apoptotic myosin cables (h) and dot plots showing the initial velocity of apoptotic cable recoil after laser microdissection (i) in control *(sqh-GFP[29B]; Dll-Gal4)* and *Arpc5* RNAi *(sqh-GFP[29B]; Dll-Gal4; UAS-Arpc5RNAi)* leg discs (n=6 and 10, respectively). Black line indicates the median. In g and i, the statistical significance was calculated using Mann-Whitney U test. ns, not significant. Scale bars are 20 μm in **a, b, e** and **f**; 2 μm in **h**.

### Leg morphogenesis occurs in a mechanically noisy environment

We next asked if folds deviate towards particular regions of the tissue in Arpc5 knockdown. We noticed that misoriented folds deviate from the predicted fold domain to more proximal or distal regions (Fig3a). Proximally, deviated folds often joined a straight cellular alignment present at the distal border of Notch activation (Fig3b, FigS3a) while distally deviated folds frequently headed towards apico-basal structures of Myosin II (Fig3d) that are not associated with apoptosis (FigS3b) and are present at various distances from the predicted fold domain. Importantly, both regions are strongly enriched in Myosin II, either apically or apico-basally, which usually coincides with high tension. Laser ablation experiments showed that tension was indeed significantly higher at the “Notch border” and at non-apoptotic apico-basal myosin structures (Fig3c, e), compared to the neighboring tissue in Arpc5 knockdown condition. Altogether, these data indicate that, when Arpc5 function is reduced, folds head towards regions of high tension. This suggests that these regions of high tension could create mechanical perturbations in close vicinity to the fold domain.

**Figure 3:**
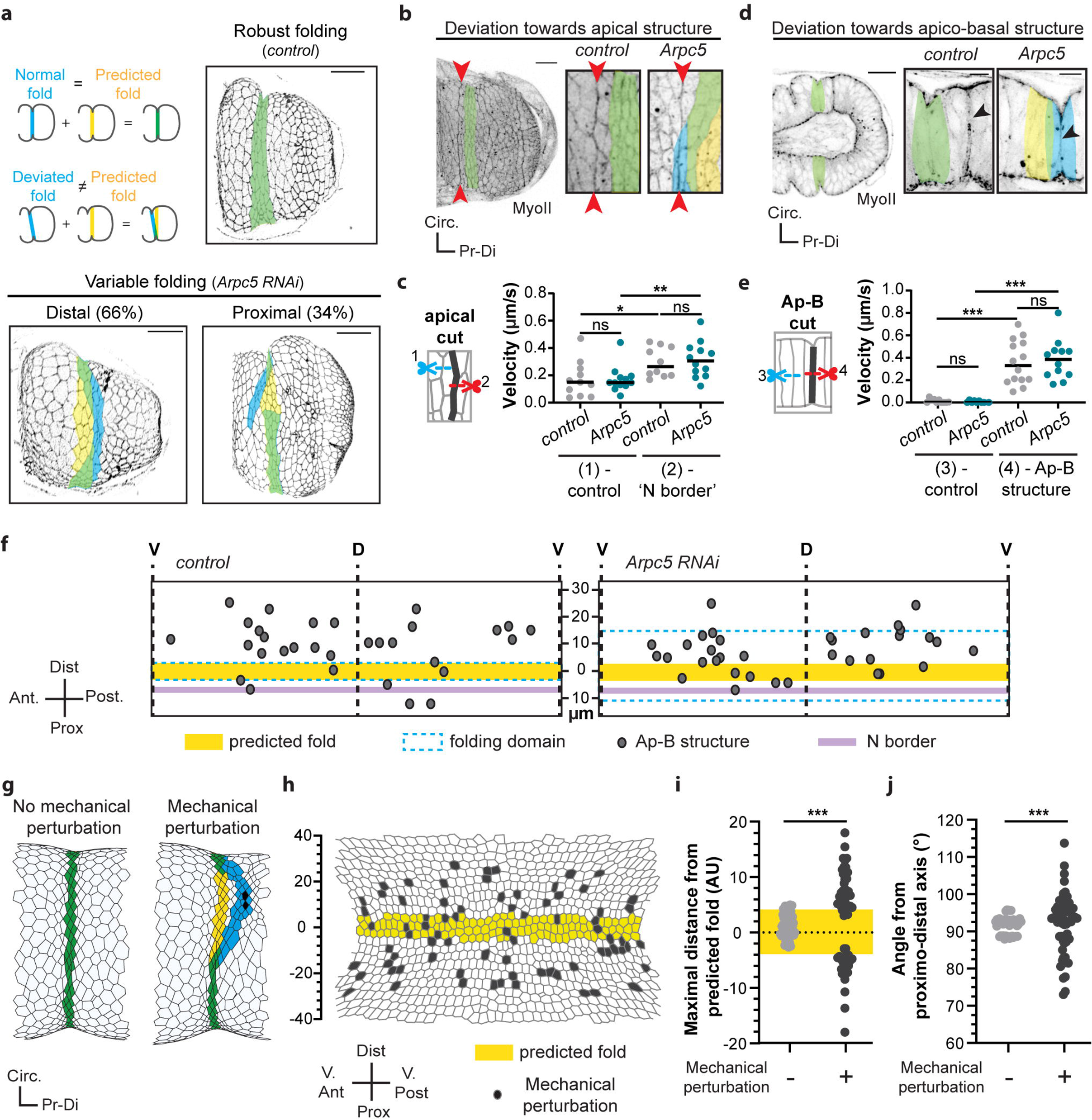
Leg morphogenesis occurs in a mechanically noisy environment (Related to FigS3, FigS4 and Movie S4). **a**, Fold morphology in *control* or *Arpc5* RNAi arm-GFP pupal leg discs and corresponding schemes. **b,d**, Confocal images showing the ‘Notch border’ (red arrowheads) **(b)** or the non-apoptotic apico-basal myosin structure (black arrowheads) **(d)** in control *(sqh-GFP[29B]; Dll-Gal4)* and Arpc5 RNAi *(sqh-GFP[29B]; Dll-Gal4; UAS-Arpc5RNAi)* conditions. **c**, Dot plots of the initial recoil velocity observed after microdissection of adherens junctions at the ‘Notch border’ or adjacent junctions in control *(sqh-RFPt[3B]; Dll-Gal4; arm-GFP)* and *Arpc5* RNAi *(sqh-RFPt[3B]; Dll-Gal4; arm-GFP, UAS-Arpc5RNAi)* leg discs (n=11, 12, 10 and 12 respectively). **e**, Dot plots of the initial recoil velocity of non-apoptotic apico-basal myosin structure or lateral membrane after laser dissection in control *(sqh-GFP[29B]; Dll-Gal4)* and *Arpc5* RNAi *(sqh-GFP[29B]; Dll-Gal4; UAS-Arpc5RNAi)* leg discs (n=14, 12, 10 and 10 respectively). In **c, e** black lines indicate the median. Statistical significance has been calculated using Mann-Whitney U test. ns, not significant; *, p-value < 0.05; **, p-value < 0.01; ***, p-value < 0.001. **f**, Rolled-out maps of the fold domain indicating the locations of mechanical perturbations observed in close vicinity of the predicted fold (yellow) in control and *Arpc5* RNAi leg discs (n=14 and 14, same genotypes as d). **g**, In silico simulations at the maximal fold depth without (left) or with (right) mechanical perturbation (challenged cells are indicated in black). **h**, Rolled-out map of mechanical perturbations random locations (grey) from 25/55 in silico simulations. 3 perturbations were integrated for each simulation. **i**, Dot plots showing the maximal distance between the real fold and the predicted fold (highlighted in yellow) in in silico simulations without (left) or with (right) mechanical perturbations. **j**, Dot plots showing the angle of the fold relative to the proximal-distal axis in in silico simulations without or with mechanical perturbations. In **a, b, d, g**, predicted fold is highlighted in yellow, real fold in blue and perfect match between them in green. In i and j, a Levene’s test was used. ***, p-value < 0.001. Scale bar represents 20 μm in a, b, d.

To figure out whether the potential mechanical interference associated with structures under high tension are either ectopically created or increased following the Arpc5 knockdown, or, alternatively, present yet masked during normal development, we analyzed the Myosin II pattern in control flies. Interestingly, both the “Notch border” and non-apoptotic apico-basal myosin structures were present in the control (Fig3b,d, FigS3) and the tension borne by these structures was comparable to that measured in Arpc5 knockdown condition (Fig3c,e, MovieS4). To further characterize tissue mechanics around the fold region, we mapped these regions of high tension and found that these mechanical perturbations are frequent and located at variable positions around the fold domain both in the control and in Arpc5 knockdown condition, suggesting the existence of a mechanical noise during fold morphogenesis (Fig3f). Altogether, these data suggest that leg fold morphogenesis is permanently challenged by surrounding remodeling events and becomes more sensitive to mechanical perturbations in the Arpc5 knockdown.

To test the impact of mechanical perturbations on fold formation, we turned to *in silico* modeling. We previously developed a vertex model able to reproduce leg fold formation both in terms of tissue shape and cellular organization (Monier et al., 2015). However, this morphogenetic event was considered as an isolated process in these simulations, and the only mechanical forces applied were those originating from the apoptotic cells located in the predicted fold domain. Therefore, we implemented the model (see M&M and FigS4) and integrated random mechanical perturbations in close vicinity to the predicted fold domain, mimicking the mechanical noise observed *in vivo* (Fig3h). Interestingly, mechanical noise appears sufficient to induce fold deviations in the simulations, mimicking the Arpc5 knockdown condition (Fig3g). We further quantified the variability of the phenotype observed when a random pattern of mechanical perturbations was integrated in the model and found that, while fold formation appeared robust in the absence of potential interferences, fold positioning and orientation become consistently more variable in the presence of mechanical perturbations in the vicinity of the predicted fold domain (Fig3i-j).

Altogether, these results strongly suggest that some kind of isolation is required between the fold domain and the neighboring tissue to avoid surrounding mechanical interference and ensure morphogenetic robustness.

### Arp2/3 controls Myosin II planar polarization

We next asked how the predicted fold domain could become insensitive to nearby mechanical noise. Strikingly, in the control we noticed that Myosin II bore a polarized junctional distribution specifically in the fold domain, with a stronger accumulation in the cellular junctions parallel to the future folds (or circumferential junctions) than in perpendicular ones (or proximal-distal junctions). This polarity, although already present at the onset of fold formation, is accentuated during fold progression and coincides with an increase of cell anisotropy (Fig4a-d). Importantly, while the total amount of Myosin II was unchanged in Arpc5 knockdown condition (FigS5a), Myosin II planar polarity was lost and cell anisotropy was no more increased in the fold domain (Fig4a-d). These observations show that folding robustness coincides with planar polarization of Myosin II in the fly leg.

**Figure 4:**
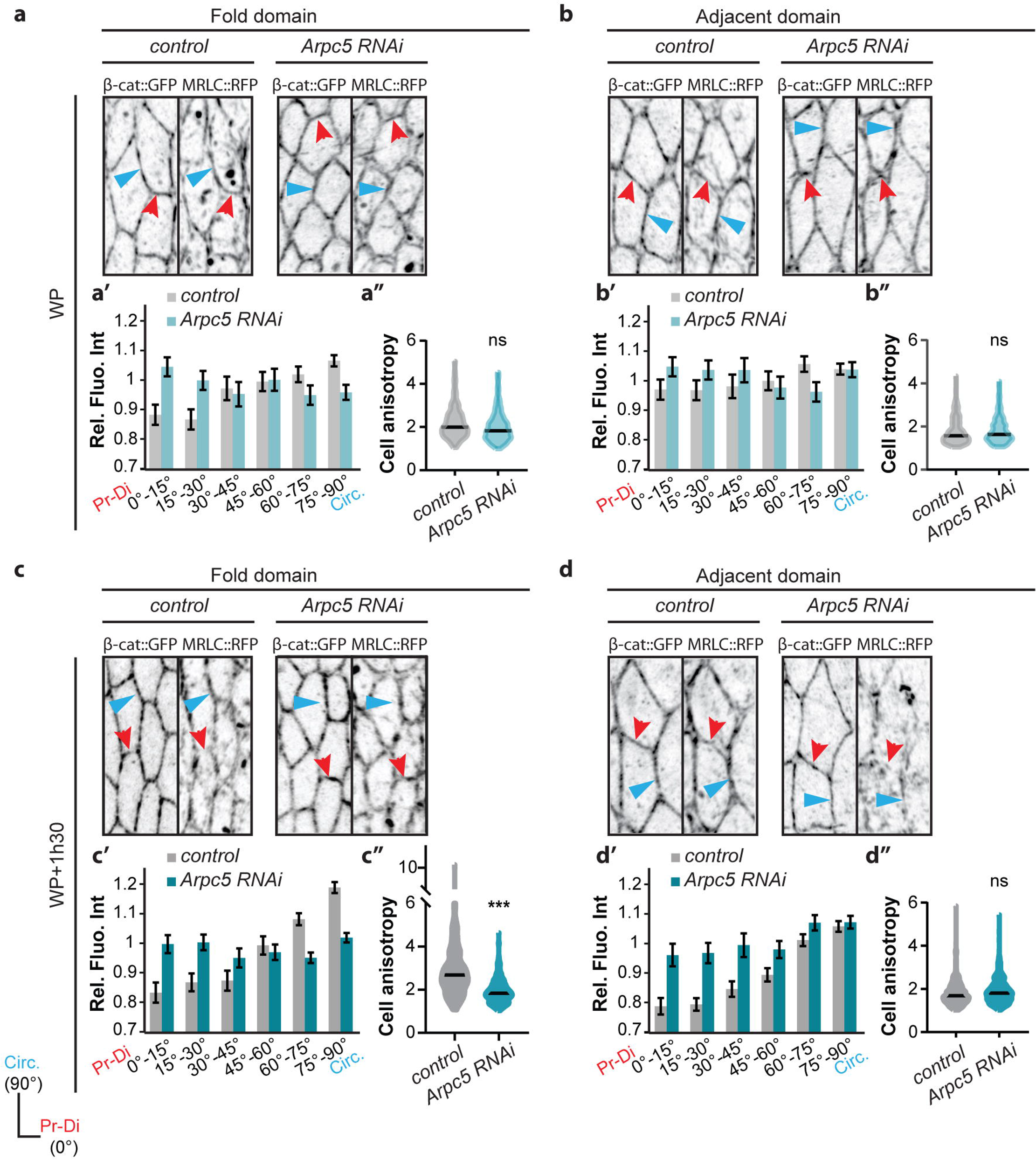
The leg epithelial tissue exhibits myosin planar polarity. (Related to FigS5) **a-d,** Close up views of confocal images showing the distribution of β-catenin-GFP and Myosin II *(sqh-RFPt[3B])* in control and *Arpc5* RNAi leg discs at the onset of fold formation (a,b) or at mid-fold stage (c,d) in the fold domain (a,c) or in the adjacent domain (b,d); **a’-d’**, Quantification of the Myosin II distribution at junctions. **a“-d”**, Quantification of the cell shape anisotropy. In **a-d**, red and blue arrowheads indicate proximal-distal and circumferential junctions, respectively. In **a’-d’**, graph bars correspond to the mean ± SEM. In **a”-d”** black lines indicate the median. Statistical significance has been calculated using Mann-Whitney U test. ns, not significant; ***, p-value < 0.001. n= 752 and n= 590 junctions in a’; n= 153 and n= 197 cells in a”; n= 603 and n= 645 junctions in b’; n= 209 and n= 254 cells in b”; n= 1107 and n= 1223 junctions in c’; n= 301 and n= 364 cells in c”; n= 1087 and n= 980 junctions in d’; n= 381 and n= 384 cells in d”.

To decipher how Arp2/3 could drive Myosin II planar polarity, we first characterized the spatial distribution of the F-actin network (using phalloidin) and of the Arp2/3 complex (using UAS-Arp3-GFP, a construct whose expression does not induce any visible defect, see FigS5b). Interestingly, we observed a polarized and anti-correlative distribution between F-actin and the Arp3-GFP fusion protein. While F-actin was preferentially accumulated in junctions parallel to the fold like Myosin II, Arp3-GFP was mainly present in junctions perpendicular to the fold (Fig5a-d). Interestingly, the polarized distribution of F-actin is lost in Arpc5 RNAi condition, indicating a role of Arp2/3 in F-actin polarity.

**Figure 5:**
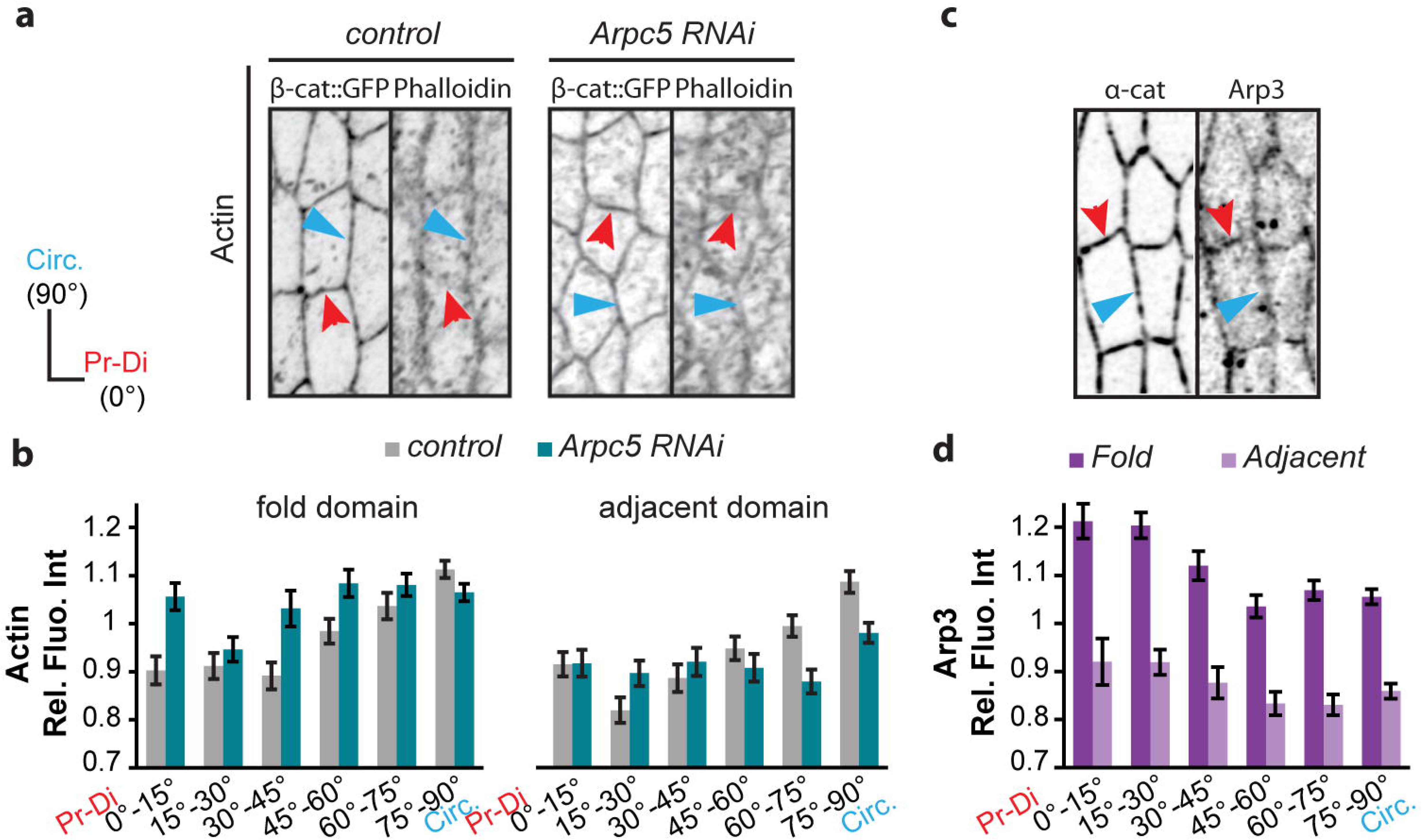
Polarized Arp2/3 drives polarized F-actin distribution in the leg epithelial tissue (Related to FigS5) **a-b**, Confocal images **(a)** showing the distribution of actin in control *(Dll-Gal4; arm-GFP)* or *Arpc5* RNAi *(Dll-Gal4; arm-GFP, UAS-Arpc5RNAi)* leg discs in the fold domain and quantification **(b)** in both the fold domain (left; n= 1004 and n= 981 junctions respectively) and the adjacent domain (right; n= 823 and n= 672 junctions respectively). **c-d**, Confocal images **(c)** showing the distribution of Arp3 *(ap-Gal4, UAS α-catenin-TagRFP, UAS Arp3-GFP)* and quantification **(d)** in the fold domain and in the adjacent domain *(n= 582 and n= 432 junctions respectively)*.

We further characterized the flows of the medio-apical Myosin II and observed a slight bias in its directionality in the control, with more movement in the proximo-distal axis than in the circumferential one (FigS5e). Interestingly this bias is lost in Arpc5 knockdown (FigS5f), suggesting that Arp2/3 could influence Myosin II polarity through the regulation of medio-apical myosin flow.

Finally, we observed a slight polarity in the distribution of adherens junctions (using β-catenin-GFP), with enrichment similar to Arp2/3. However, this polarity is unperturbed in Arpc5 knockdown (FigS5c-d).

### Planar polarity ensures morphogenetic robustness

To test if perturbing Myosin II planar polarity was sufficient to alter fold robustness *in vivo*, we decided to disrupt Myosin II polarity by an independent method. We generated a new variant of the nanobody-based GFP-trap technique previously described (Harmansa et al., 2017; Harmansa et al., 2015), which traps the endogenous Myosin II-GFP fusion protein (Myosin Regulatory Light Chain, MRLC-GFP) at adherens junctions (Fig6a-b, FigS6a, see M&M). We found that the distribution of junctional Myosin II is mainly isotropic in this condition, although not totally homogeneous. Interestingly, using conditional expression of this AJs-GFP-trap construct in the presence of Myosin II-GFP, we frequently observed deviated folds, mimicking the defects observed in the Arpc5 knockdown (Fig6c), while expression of AJs-GFP-trap in the absence of any GFP does not alter leg morphogenesis (FigS6b). We also observed that the variability of fold positioning, fold orientation and fold parallelism was significantly increased in this condition of Myosin II polarization defect compared to the control (Fig6d-f). These experiments show that decreasing or abolishing Myosin II planar polarity through two independent strategies (Arpc5 knockdown and AJs-GFP-trap/Myosin II-GFP) leads to an increase of variability of fold orientation in the fly leg.

**Figure 6:**
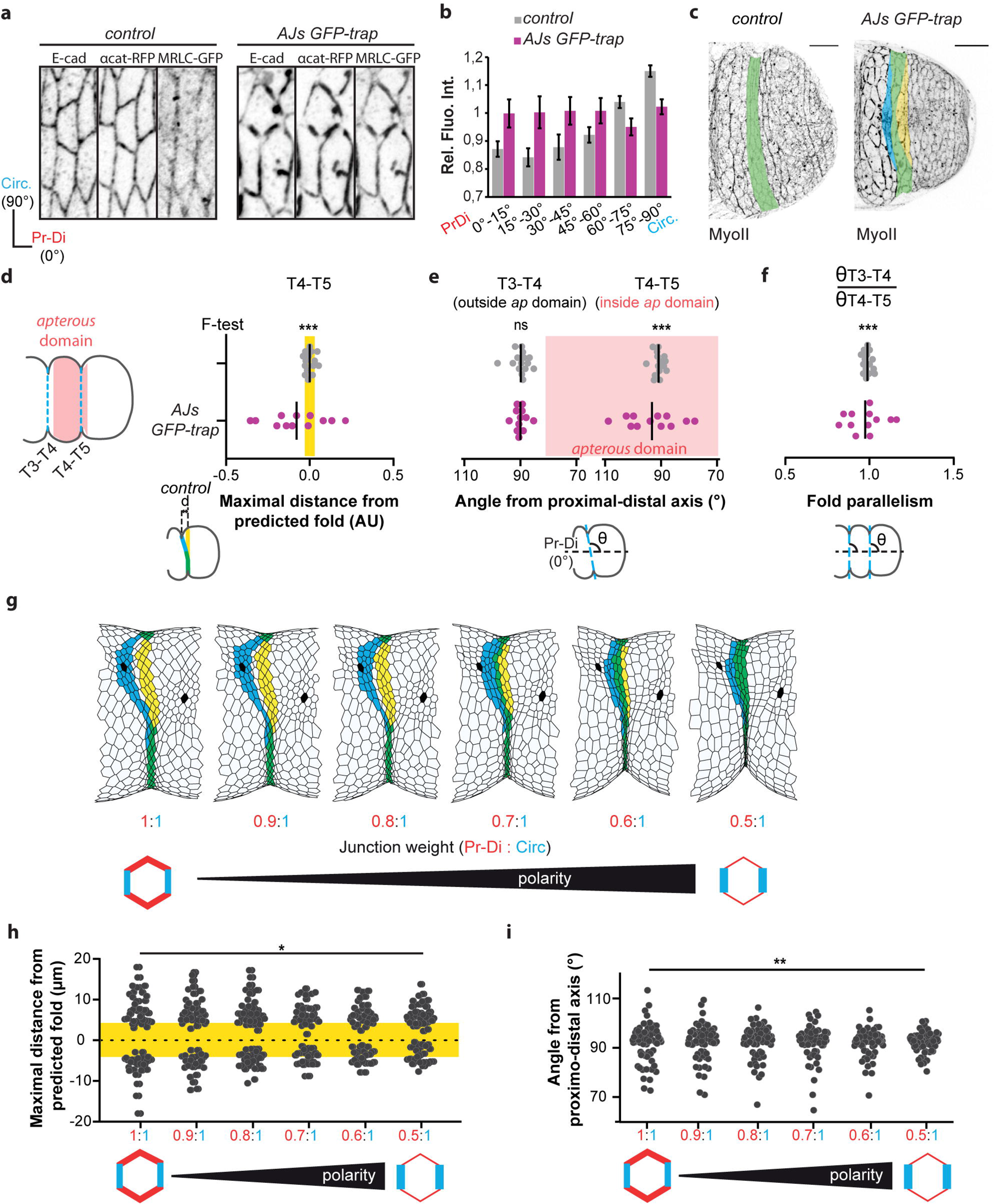
Planar polarity is required for morphogenetic robustness (Related to FigS6 and FigS7). **a**, Confocal images showing the distribution of E-cadherin, α-catenin and Myosin II in control *(sqh-GFP[29B]; tubG80ts; ap-Gal4, UAS α-catenin-TagRFP)* or AJs GFP-trap *(sqh-GFP[29B]; tubG80ts; ap-Gal4, UAS vhhGFP4-α-catenin-TagRFP)* leg discs. **b**, Quantification of Myosin II distribution at junctions in leg discs in both conditions (n= 1009 and n= 564 junctions respectively). **c**, Confocal images showing fold morphology in control or AJs GFP-trap leg discs (deviated fold represents 0/15 and 8/12, respectively). Predicted fold domain is highlighted in yellow, real fold in blue and perfect match between them in green. Scale bar represents 20 μm. **d**, Dot plots showing the relative maximal distance between the real fold and the predicted fold (highlighted in yellow) in control or AJs GFP-trap leg discs (n=15 and 12 legs, respectively). **e**, Dot plots showing the angle of T3-T4 and T4-T5 folds relative to the proximal-distal axis in control or AJs GFP-trap leg discs (n=15 and 12 legs, respectively). **f**, Dot plots showing the fold parallelism in control or AJs GFP-trap leg discs (n=15 and 12 legs, respectively). in **d-f**, A F-test of equality of variances has been used to compare the phenotypic variances. ***, p-value < 0.001. **g**, In silico simulations including mechanical perturbations (challenging cells are shown in black) for different values of junction weight ratio (i.e. tension anisotropy). Predicted fold domain is highlighted in yellow, real fold in blue and perfect match between them in green. **h**, Dot plots showing the maximal distance between the real fold and the predicted fold (highlighted in yellow) for different values of junction weight ratio (polarity). (n= 55 simulations in each cases). **i**, Dot plots showing the angle of the fold relative to the proximal-distal axis for different values of junction weight ratio (polarity). (n= 55 simulations in each cases). In h and i, a Levene’s test has been used to compare the variance of phenotypes. *, p-value < 0.05; **, p-value < 0.01.

We then wondered whether restoring Myosin II polarity in a non-polarized tissue could rescue the defects of fold deviation. Although not feasible *in vivo*, we could address this question using *in silico* modeling, asking whether fold deviation caused by local mechanical perturbations (see Fig3g) could be rescued by the introduction of planar polarized junctional Myosin II. In order to integrate Myosin II planar polarity in the model, we first checked if tension pattern could be inferred by myosin distribution in this tissue. Using laser ablation, we found that circumferential junctions bear more tension than proximal-distal junctions in the control, while tension appears independent of junction orientation in Arpc5 knockdown (FigS7a-b), consistent with the respective distributions of Myosin II observed in these conditions. We then mimicked Myosin II planar polarity and the associated tension anisotropy by the attribution of different values of junctional tension depending on junction orientation in our model (see M&M and FigS7c-e). Fold robustness was unaffected by the integration of tension polarity in the model in the absence of external perturbations (FigS7f), while interestingly, gradual increase of tissue polarity favors fold straightness and insensitivity to surrounding perturbations (Fig6g). We further quantified fold morphogenetic robustness in our theoretical model for different degrees of polarity. Interestingly, increasing polarity decreases the degree of deviation of the fold and thus decreases the variability in fold directionality (Fig6h-i and FigS7g-h). Altogether, *in vivo* manipulations and *in silico* modeling indicate that the planar polarization of tissue tension favors mechanical isolation of fold formation, which ultimately ensures morphogenetic robustness.

### Planar polarity favors directional force transmission

We then asked how planar polarity of Myosin II and the associated polarized tension could protect morphogenesis from surrounding mechanical disturbances, ensuring robust fold formation.

We first performed laser ablation at the level of a tricellular junction in the fold domain and followed the recoil of vertices in the neighboring tissue, along several cell diameters (Fig7a). We observed stronger recoil in the direction of the fold in the control (Fig7b-c). On the contrary, in the Arpc5 knockdown condition, recoil of vertices up to 3-4 cell diameters becomes similar irrespective of the direction (Fig7d-e). This experiment reveals the existence of an anisotropic multicellular mechanical coupling, which results in tissue-scale tension anisotropy.

**Figure 7.**
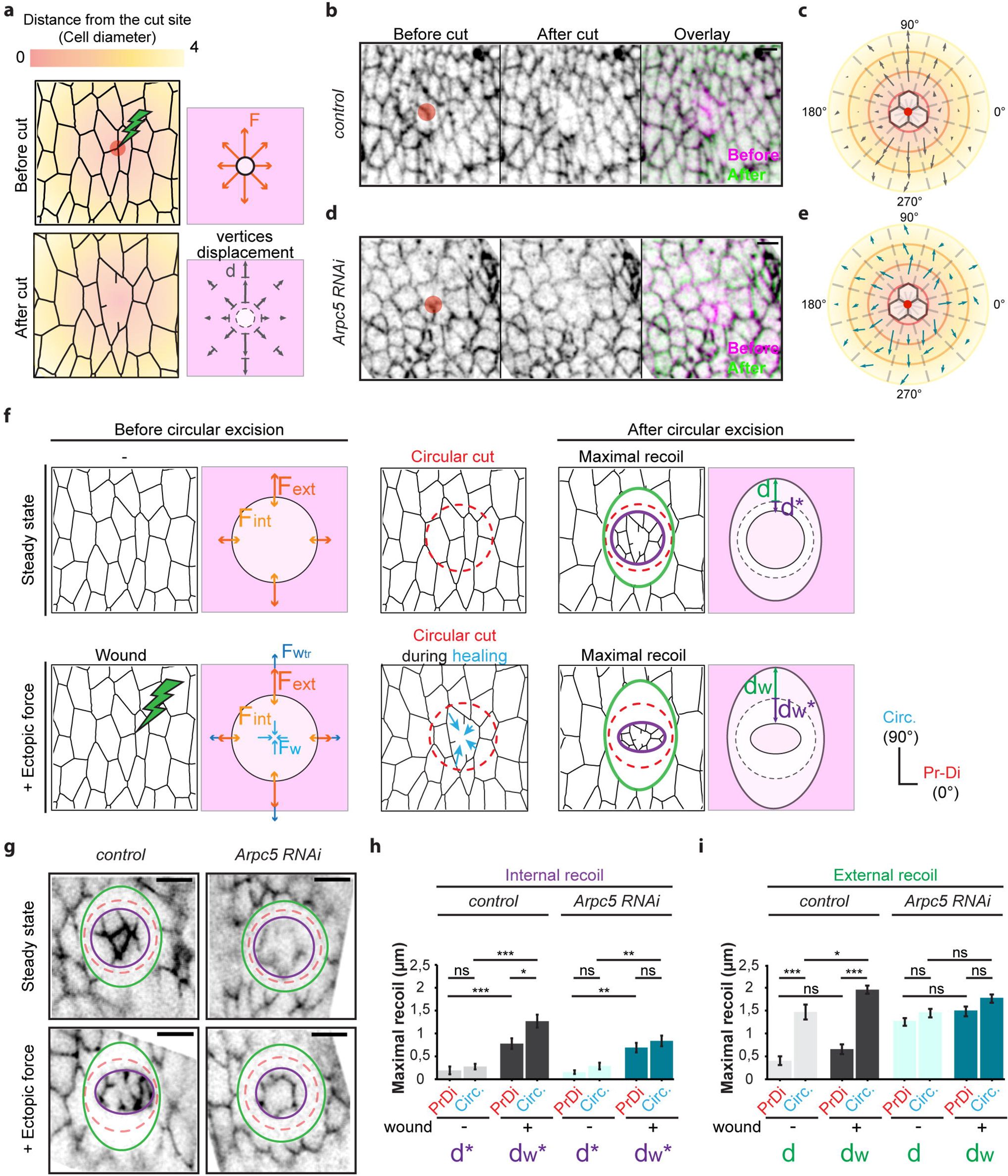
Planar polarity favors force channeling. **a**, Schematics showing the impact of the wound (left), the forces in presence (F), and the maximum displacement of vertices at various cell diameters (d). **b,d**, Z-projections of control (Dll-Gal4; arm-GFP) and *Arpc5* RNAi *(Dll-Gal4; arm-GFP, UAS-Arpc5RNAi)* leg discs before (left) and after (middle) the cut. Overlay are shown on the right. Red circles indicate site of ablation. Scale bar represents 2 μm. **c,e**, Average displacement of vertices (represented by vectors) at different distances and angles from the cut site (center) in control **(b)** and Arpc5 RNAi (c) (n= 8, 10 legs respectively). Distance from the cut is color coded as in a. **f**, Schematics indicating the absence (top) or the presence (bottom) of wound and the forces in presence at the level of the circular cut (left) and the external and internal maximal recoils after circular ablation (right). In the absence of wound (top) the internal release d* depends on the residual stress present in the circular isolated tissue fragment, while the external release d depends on the tension present in the rest of the tissue. In the presence of wound-healing traction force (bottom), the internal release dw* depends on both the residual stress present in the circular tissue fragment (Fint) plus the wound-healing traction force (Fw), while the external release dw depends on both residual stress in the whole tissue plus the force transmitted in response to the wound-healing traction force (Fwtr). **g**, Confocal images of the control and Arpc5 RNAi leg discs (same genotypes as in b). External and internal maximal recoils of the tissue are indicated by the green and purple lines respectively, while the initial positioning of the circular ablation is indicated by a dashed red line. Scale bar represents 5 μm. **h-i**, Quantifications of internal (h) and external (i) maximal recoils of the tissue in the absence (light grey and light blue) or the presence (dark grey and dark blue) of wound in control and Arpc5 RNAi leg discs (n= 10, 10, 12, 10 respectively). Statistical significance has been calculated using Mann-Whitney U test. ns, not significant; *, p-value < 0.05; **, p-value < 0.01; *** p-value < 0.001. Graph bars correspond to the mean ± SEM.

We hypothesized that Myosin II polarized distribution could generate a bias in long-range force transmission, avoiding force scattering across the tissue. To specifically test how discrete forces, generated locally, are transmitted along the tissue, we used wound healing as a way to generate local forces in a spatio-temporally controlled manner (Fernandez-Gonzalez and Zallen, 2013). We next analyzed the propagation of forces across the tissue. We wounded the tissue locally by laser ablation at the level of a tricellular junction, waited for the tissue to repair, and as soon as the healing started generating local forces, performed a circular cut around the healing region and observed the recoil (Fig7f-g, + ectopic force, bottom). A similar experiment in the absence of a preliminary wound gave access to the steady state pattern of tensions within the leg tissue (Fig7f-g, steady state, top row). Thus, in the absence of wound, the internal release (d*) depends on the tension, or residual stress, present in the circular isolated tissue fragment, while the external release (d) depends on the tension present in the rest of the tissue. In the presence of wound-healing traction force, the internal release dw* mostly depends on both the residual stress present in the circular tissue fragment (Fint) plus the wound-healing traction force (Fw), while the external release dw mostly depends on both residual stress in the whole tissue (Fext) plus force transmission (FWtr). This allows us to estimate the wound-healing traction force by comparing dw* and d*, while we can estimate the transmission of forces created in response to healing by comparing dw and d.

We first analyzed internal recoil and observed that while the recoil is isotropic in the absence of Myosin II planar polarity (compare dw*_PrDi_ and dw*_Circ_ in Arpc5 RNAi, Fig7h), it is anisotropic in the presence of Myosin II planar polarity (compare dw*_PrDi_ and dw*_Circ_ in control, Fig7h). This anisotropy suggests that the traction force due to the healing (Fw) could be anisotropic in the control. Regarding the external recoil, without wound healing, circular cutting induced an ovoid-shaped recoil, indicating that tissue tension was stronger in the circumferential axis in the wildtype condition, while it results in circular recoil in Arpc5 RNAi condition, indicating that tension anisotropy was lost (compare d_PrDi_ to d_Circ_ in control and d_PrDi_ and d_Circ_ in Arpc5 RNAi in Fig7i). In the presence of wound healing, the tendency is the same (compare d_WPrDi_ and d_WCirc_ in control and d_WPrDi_ and d_WCirc_ in Arpc5 RNAi in Fig7i). Interestingly, while an increase in internal recoil is visible both in proximo-distal and circumferential axis in the presence of wound (compare d*_PrDi_ with dw*_PrDi_ and d*_Circ_ with dw*_Circ_ in the control and Arpc5 RNAi in Fig7h), the external recoil is specifically accentuated along the circumferencial axis in the presence of Myosin II planar polarity (compare d_Circ_ with dw_Circ_ and d_PrDi_ with dw_PrDi_ in control, Fig7i). Thus, even if initial wound healing traction force might be anisotropic, these experiments allow us to propose that in the presence of Myosin II planar polarity, forces (both traction forces and forces transmitted in response) are transmitted preferentially in the circumferential direction (in dark grey in Fig7h-i). On the contrary, in the absence of Myosin II planar polarity, no significant increase in external recoil is observed (compare d_Circ_ and dw_Circ_ as well as d_PrDi_ and dw_PrDi_ in Arpc5 RNAi, Fig7i), showing that forces might spread homogeneously around the wound healing region in this condition (in dark blue in Fig7h-i).

Altogether, these experiments indicate that forces are not homogeneously transmitted in the junctional plane of the tissue but are rather channeled in the presence of Myosin II planar polarity.

## Discussion

Our study provides direct evidence of a mechanism protecting morphogenesis from environmental perturbations, ensuring tissue shape robustness on top of developmental patterning through the channeling of mechanical forces. This mechanism involves Arp2/3 complex, which controls junctional tension anisotropy through Myosin II planar polarity, and avoid apical force scattering, thus favoring force transmission along the circumferential axis. This ultimately provides resistance to mechanical perturbations that appear randomly in the surrounding tissue and create a mechanically noisy environment.

Our work reveals that Arp2/3 complex plays a crucial role in establishing a planar polarized distribution of Myosin II in the developing leg tissue. We explored several hypotheses to explain the involvement of Arp2/3 in Myosin II planar polarity. Arp2/3 has been related to E-cadherin endocytosis (Georgiou et al., 2008). By favoring E-cadherin endocytosis at junctions, it could indirectly impact Myosin II polarity. However, adherens junction polarity is not perturbed in Arp2/3 knockdown, ruling out this hypothesis. Myosin II planar polarity in the embryo depends on medial-apical flows of acto-myosin (Rauzi et al., 2010). Consistently, we observed a slight bias in the directionality of medial-apical myosin flow in the control that is lost in Arpc5 knockdown. This indicates that Arp2/3 influences Myosin flow, however, these results are complex and require further investigation. Finally, Arp2/3, which is involved in the formation of branched actin network, is enriched in the junctions that are perpendicular to the fold, while F-actin is more abundant in circumferential junctions. This suggests that the distribution and the density of F-actin networks is itself polarized and that proximo-distal junctions are enriched in branched actin, while circumferential junctions would be enriched in linear actin. Since Myosin II has been observed to preferentially accumulate with linear actin network (Michelot and Drubin, 2011), this polarity of the actin networks would favor the accumulation of Myosin II along circumferential junctions and thus drive Myosin II planar polarity.

This work also highlights that Myosin II planar polarity could protect a morphogenetic process from the surrounding noise, by playing a role in long-range force chanelling and forming a sort of mechanical fence. Myosin II planar polarity has been described in different contexts: germ band extension in Drosophila embryo where it drives tissue elongation (Bertet et al., 2004), but also neural tube closure in chicken (Nishimura et al., 2012), where it favors polarized junction shortening and tissue folding, or at compartment boundaries in Drosophila and zebrafish where it maintains a straight borders (Calzolari et al., 2014; Major and Irvine, 2006; Monier et al., 2010). However, its role in the protection of morphogenesis from external perturbations was unexpected. Indeed, so far, most of the studies related to morphogenetic robustness converge on the regulation of gene expression patterns in response to morphogen gradients and consider acto-myosin either as a component of the “core toolbox” responsible for building a new shape or as a mechanism to buffer local heterogeneity in positional information by exerting a feedback on cell-fate decisions (Gilmour et al., 2017). However, the influence of tissue mechanics in maintaining low variability of a specific shape when challenged by intrinsic perturbations in close proximity was unexplored.

Interestingly, two recent papers addressed the robustness of tissue invagination. While the work of Yevick et al highlights the importance of mechanical redundancy to resist to accidental damage (Yevick et al., 2019), Eritano et al reveal the role of mechanical coupling as an intrinsic property of morphogenesis to buffer small variability in gene expression patterns (Eritano et al., 2020). The present study appears complementary to these previous works, showing how morphogenesis is naturally protected from mechanical perturbations occurring randomly in the surroundings, by creating a fence through Arp2/3-dependent junctional myosin II planar polarity. It further reveals that tissue mechanics not only buffer genetic information but can take over the positional information given by patterning genes since folds can considerably deviate from the pre-established program if forces are not properly channeled.

Finally, this work further reveals that fold morphogenesis in the Drosophila developing leg occurs in the presence of mechanical noise, as shown by mechanical perturbations randomly distributed in the developing tissue at the time of fold formation. Since the occurrence of several developmental processes in the same time window is frequently observed during development, it is tempting to speculate that mechanical noise could be a general feature of morphogenesis and that mechanical isolation could be required in a wide variety of morphogenetic processes to avoid force scattering and maintain morphogenetic robustness. Thus, this process of force channeling through Myosin II polarized distribution could be a general way to isolate a particular morphogenetic process from surrounding events, preventing any interference between closely located events and favoring robustness.

## Acknowledgements

We thank Michel Labouesse and Yohanns Bellaiche for their constructive comments on the manuscript. MS’s lab is supported by grants from the European Research Council (ERC) under the European Union Horizon 2020 research and innovation program (grant number EPAF: 648001), from the Institut National de la Santé et de la Recherche Médicale (Inserm, Plan cancer 2014-2019) and from the association Toulouse Cancer Santé (TCS, ApoMacImaging: 171441). EM has a post doc fellowship from the Association pour la Recherche contre le Cancer (ARC). ST has a CIFRE fellowship from the ANRT.

## Author contributions

EM conceived and performed the experiments in fly leg discs. ST participated in modeling conception and realized all the simulations. EM and ST analyzed and quantified the data. BM initiated the project, participated in experimental conception and article writing. GG supervised the modeling conception. MS supervised the project, wrote the paper and provided the funding. CR helped with the analysis of Rpr and Dpn pattern, and PIV analysis.

## Competing interest declaration

The authors declare no competing financial interests.

## Corresponding authors

Correspondence and requests should be addressed to M.S. for biology and materials and to G.G. for modeling.

## Star Method

### Key Resources Table

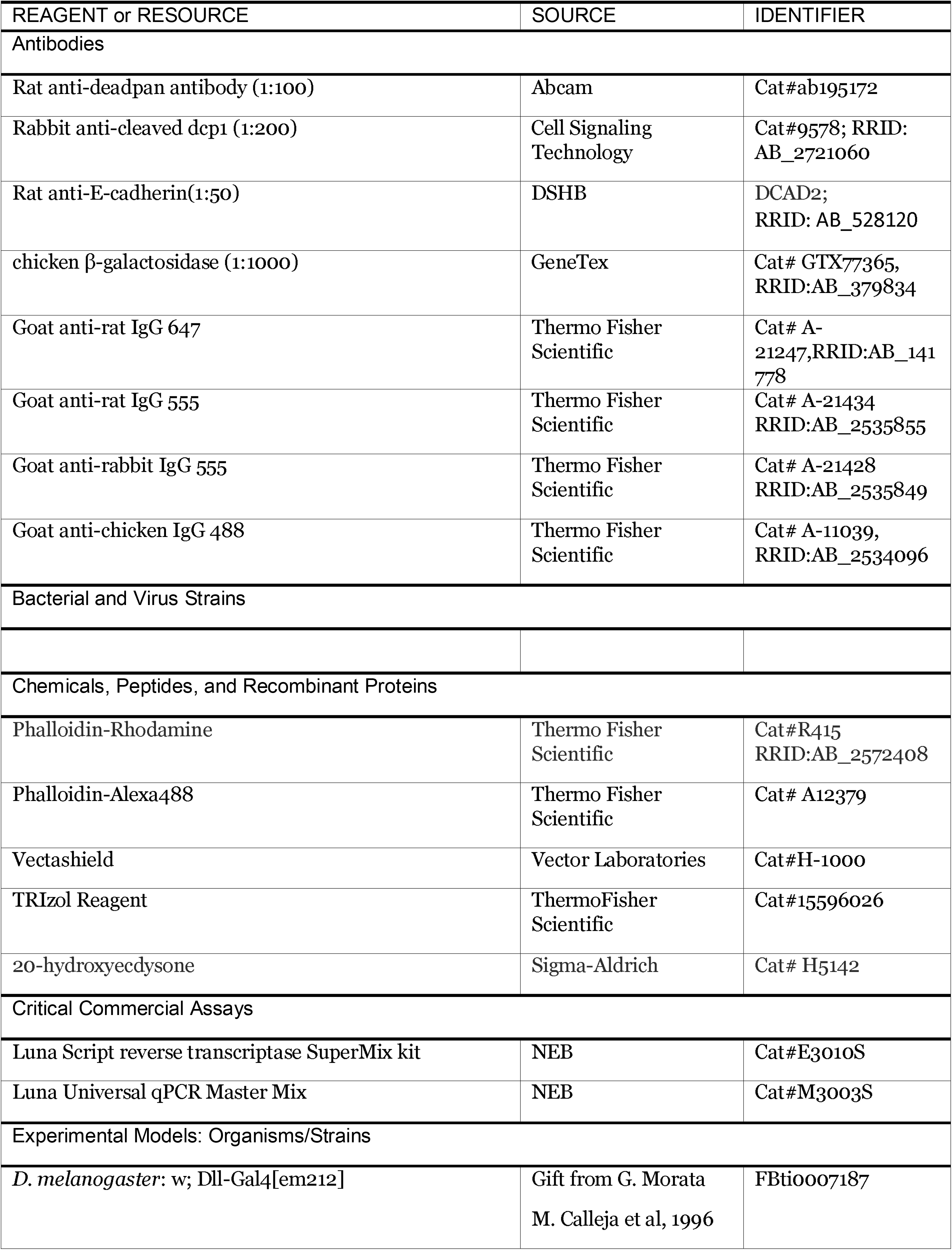
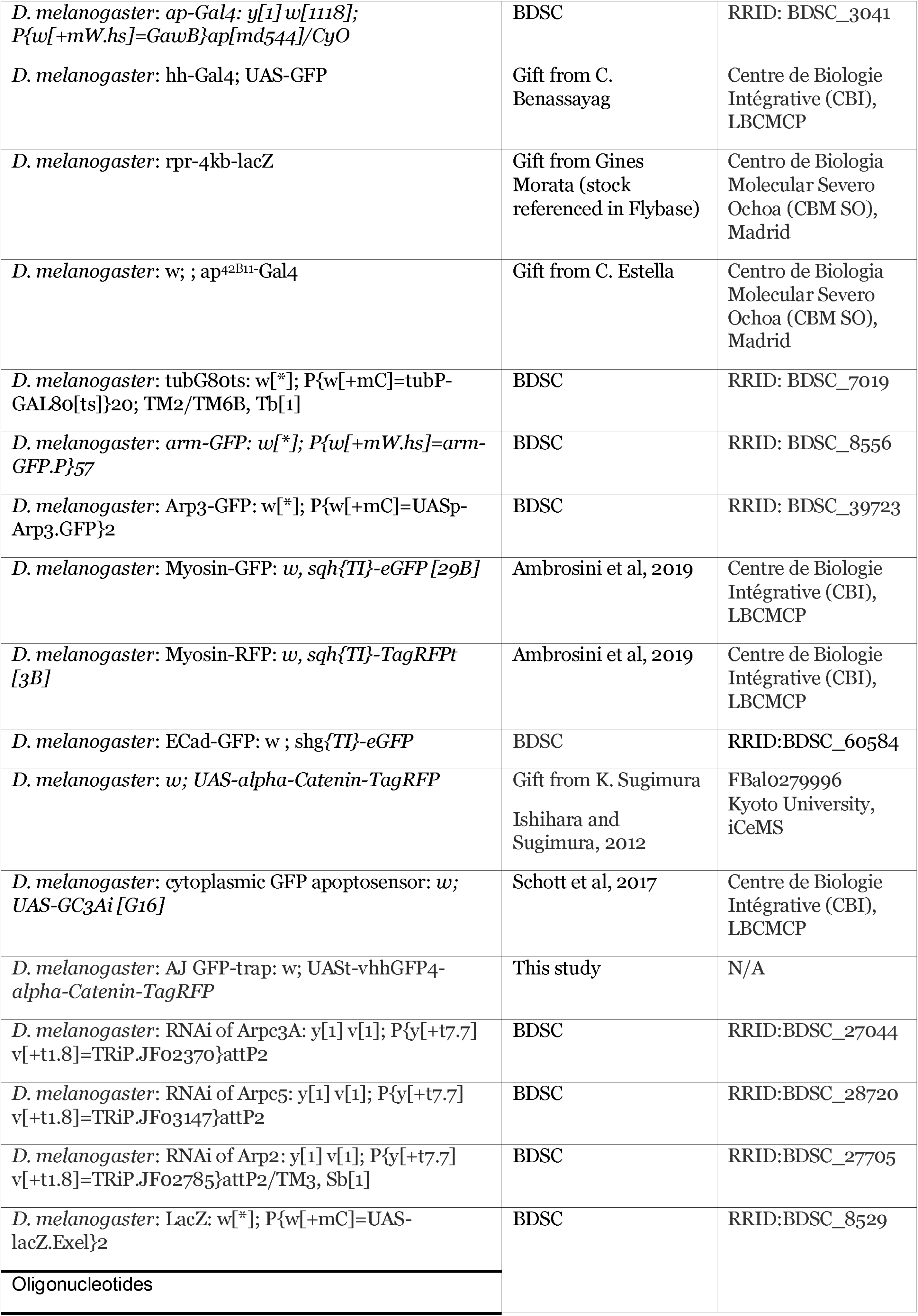
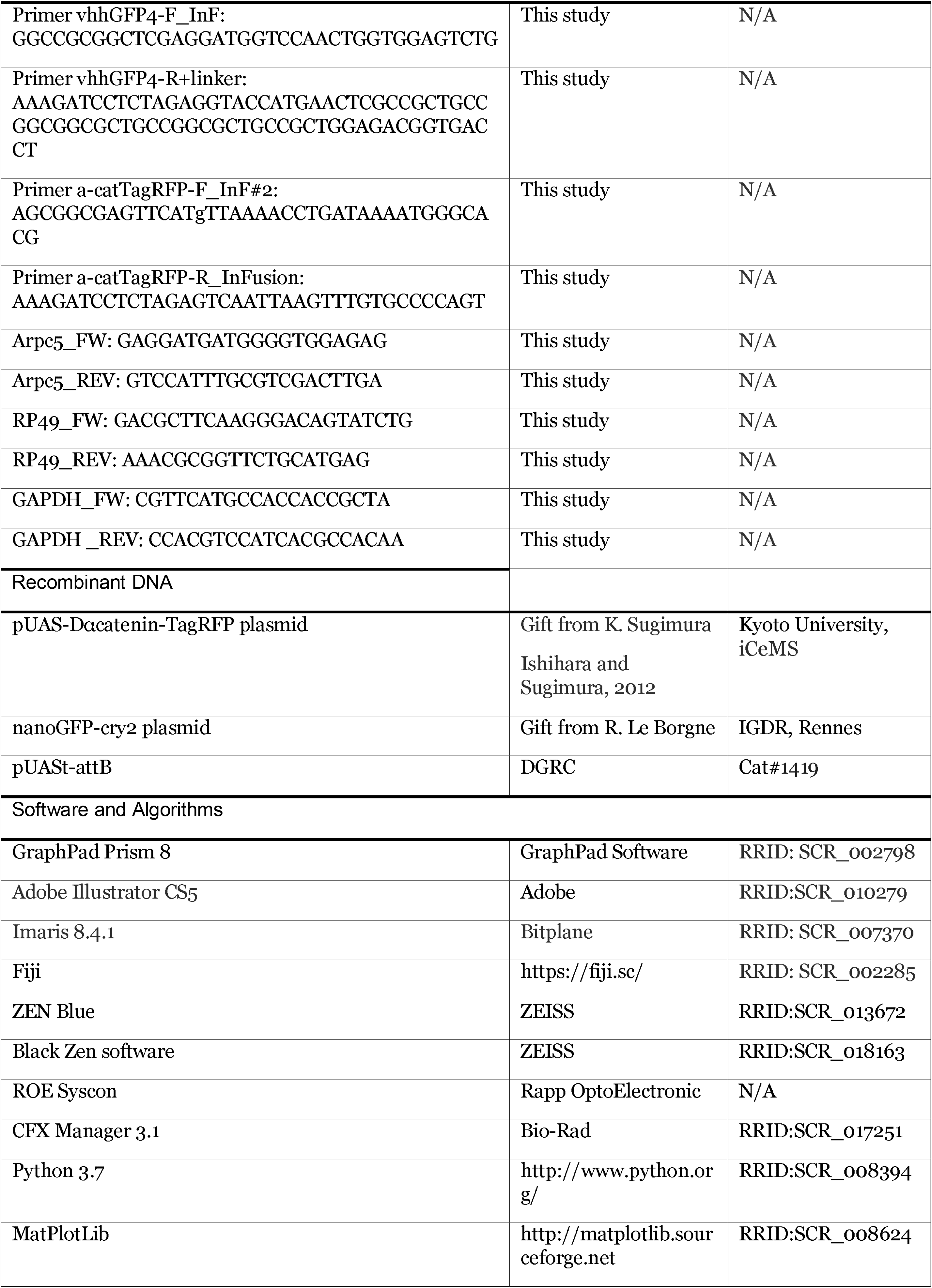
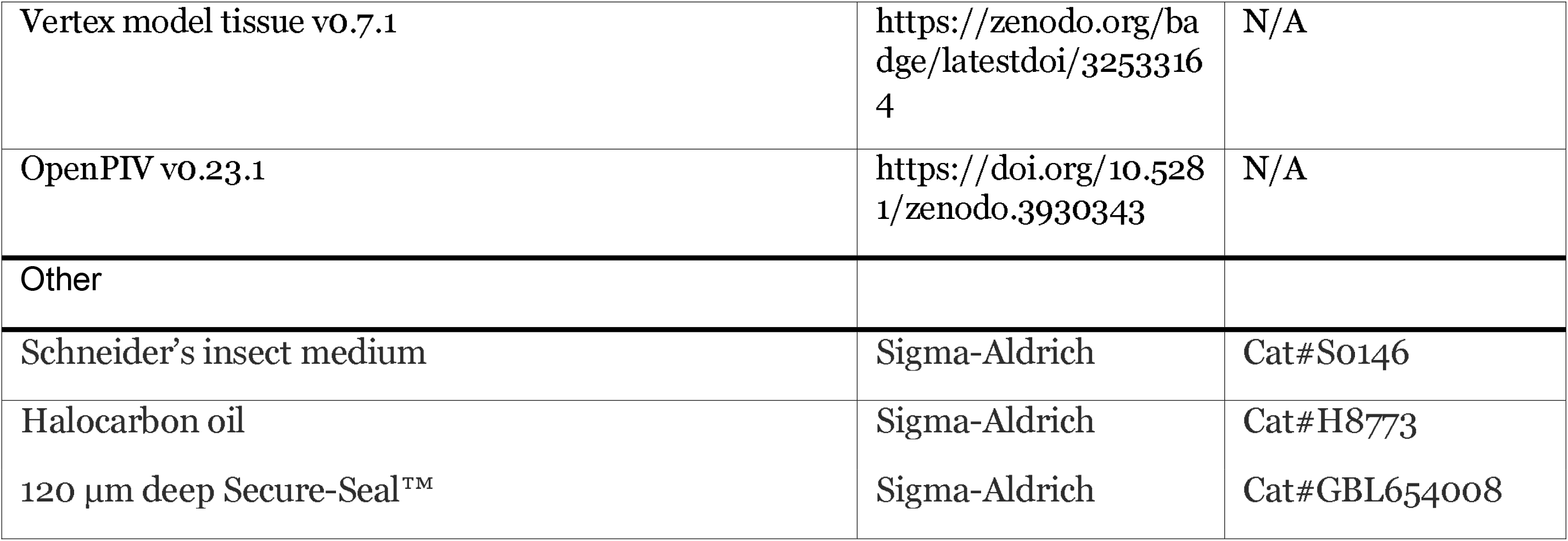

### Contact for Reagent and Resource Sharing

Further information and requests for resources and reagents should be directed to and will be fulfilled by the Lead Contact, Magali Suzanne.

### Experimental model and subject details

#### Experimental Animals

The animal model used here is Drosophila melanogaster, in a context of in vivo/ex vivo experiments. In order to respect ethic principles, animals were anesthetized with CO2 (adults) before any manipulation. To avoid any release of flies outside the laboratory, dead flies were frozen before throwing them. Stocks of living flies were conserved in incubators, either at 18 or 25 degrees to maintain the flies in optimal condition. Genotypes and developmental stages are indicated below. Experiments were performed in both males and females indifferently. Loss of function experiments using RNAi were carried out at 30 degrees.

#### Drosophila melanogaster

ap^md544^-Gal4, arm-arm-GFP, tubG80ts, UAS-Arp3-GFP and E-Cad-GFP knock-in were obtained from Bloomington Drosophila Stock Center (BDSC). sqh-eGFP^KI^[29B], sqh-RFPt^KI^[3B] and the GFP apoptosensor UAS-GC3Ai were described previously (Ambrosini et al., 2019; Schott et al., 2017). hh-Gal4; UAS-GFP is a gift from C. Benassayag. Dll^EM212^-Gal4 and rpr-4kb-lacZ are gifts from G. Morata. ap^42B11^-Gal4 and UAS-α-catenin-TagRFP are gifts from C. Estella and K. Sugimura.

RNAi experiments were realized using UAS-Arpc3A-RNAi (JF02370), UAS-Arpc5-RNAi (JF03147), UAS-Arp2 RNAi (JF02785), UAS-lacZ, obtained from BDSC.

### Method details

#### AJ GFP-trap construct

To trap Myosin-GFP at junctions, we constructed a p*UASt–vhhGFP4-*α*-catenin-TagRFP* (AJs GFP-trap). To do so, α*-catenin-TagRFP* was amplified by PCR with specific primers from a pUAS-Dαcatenin-TagRFP (kindly provided by Dr K. Sugimura). vhhGFP4 was amplified by PCR from nanoGFP-cry2 plasmid (gift from Dr Le Borgne). A 36nt sequence was added in the reverse primer to create a GSAGSAAGSGEF linker between the GFP-trap sequence and α-catenin-TagRFP sequence. These two fragments were successively cloned using InFusion technology in pUASt-attB plasmid cut with KpnI.

The resulting p*UASt–vhhGFP4-*α*-catenin-TagRFP* plasmid injection was performed by the CBMSO Drosophila Transgenesis Service (Madrid, Spain) using flies carrying attP2 landing sites on the third chromosome to produce transgenic flies.

#### RNA and RT-qPCR analyses

RNAi efficiency was assessed by mRNA phenol/chloroform extraction and reverse transcription was done using the Luna Script reverse transcriptase SuperMix kit (NEB – E3010S). cDNAs were quantified by qPCR using the Luna Universal qPCR Master Mix (NEB – M3003S) (primer sequences: Arpc5_FW GAGGATGATGGGGTGGAGAG and Arpc5_REV GTCCATTTGCGTCGACTTGA) and normalized against RP49 and GAPDH cDNA levels (primer sequences: RP49_FW GACGCTTCAAGGGACAGTATCTG; RP49_REV AAACGCGGTTCTGCATGAG and GAPDH_FW CGTTCATGCCACCACCGCTA; GAPDH _REV CCACGTCCATCACGCCACAA). The relative normalized cDNA expression levels were calculated using the DeltaDeltaCt method from Bio-Rad CFX Manager 3.1 software.

#### Immunofluorescence

Imaginal leg discs were dissected after 48h at 29°C at prepupal stage (0, 1.5 or 2.5 hours after pupae formation – APF) in PBS 1X. Imaginal discs were fixed 20’ in paraformaldehyde 4% diluted in PBS 1X, then washed in PBS 1X and either mounted in Vectashield (Vectors laboratories) or extensively washed in PBS-Triton 0.3%-BSA 1% (BBT) and incubated overnight at 4°C with appropriate dilutions of primary antibodies in BBT. Rat anti-deadpan antibody (Abcam – ab195172) was used at 1:100 dilution, rabbit anti-dcp1 (Cell Signaling Technology – 9578S) at 1:200, rat anti-E-cadherin antibody (Developmental Studies Hybridoma Bank – DCAD2) at 1:50 and chicken β-galactosidase (GeneTex – GTX77365) at 1:1000. After washes in BBT, imaginal leg discs were incubated at room temperature for 2 h with 1:200 anti-rat IgG 647, anti-rat IgG 555, anti-rabbit IgG 555 or anti-chicken IgG 488 (obtained from Interchim) with or without phalloidin (Alexa488 at 1:500 or rhodamin at 1:500 – Fisher Scientific). Then, samples were washed in PBS-Triton 0.3%, suspended in Vectashield (Vectors laboratories) and mounted on slides.

#### Ex vivo culture of leg imaginal disc

Imaginal leg discs were dissected from prepupal stage (1.5 hours APF at 29°C) in Schneider’s insect medium (Sigma-Aldrich) supplemented with 15 % fetal calf serum and 0.5 % penicillin-streptomycin as well as 20-hydroxyecdysone at 2 μg/mL (Sigma-Aldrich, H5142). Leg discs were transferred on a slide in 12 μL of this medium in a well formed by a 120 μm-deep double-sided adhesive spacer (Secure-SealTM from Sigma-Aldrich). A coverslip was then placed on top of the spacer. Halocarbon oil was added on the sides of the spacer to prevent dehydration.

#### Confocal imaging

Samples were analyzed using a LSM880 confocal microscope fitted with a Fast Airyscan module (Carl Zeiss) and equipped with a Plan-Apochromat 40x/NA 1.3 Oil DIC UV-IR M27 objective. Z-stacks were acquired using either the laser scanning confocal mode or the High Resolution mode (Airyscan) with a pixel size of 0.046 μm/pixel and a *z*-step of 0.220 um. Airyscan Z-stacks were processed in ZEN software using the automatic strength (6 by default) and the 3D method.

#### Laser ablation

Laser ablation experiments were performed using a pulsed DPSS laser (532 nm, pulse length 1.5 ns, repetition rate up to 1 kHz, 3.5 μJ/pulse) steered by a galvanometer-based laser scanning device (DPSS-532 and UGA-42, from Rapp OptoElectronic, Hamburg, Germany) and mounted on a LSM880 confocal microscope (Carl Zeiss) equipped with a 63x C-Apochromat NA 1.2 Water Corr objective (Carl Zeiss). Photo-ablation of apical junction was done in the focal plane by illuminating at 70 % laser power during 1 s. This focal plane was acquired every 0.551 s, during 10 s before and at least 45 s after ablation, with a pixel size of 0.13 μm/pixel. Photo-ablation of apico-basal Myosin II enrichment was done in the focal plane by illuminating at 100 % laser power during 2-2.5 s along a 45-50 pixels line. This focal plane was acquired every 0.551 s, during 5 s before and at least 45 s after ablation, with a pixel size of 0.13 μm/pixel. Data analysis was performed with the ImageJ software using a homemade macro.

For experiments on tissue scale tension anisotropy (Fig7a-e), a Z-stack of 7 slices was acquired every 1.774s during 10s before and 3 minutes after ablation, with a pixel size of 0.13 μm/pixel. Photo-ablation of a vertex was done during the stack acquisition by illuminating at 75 % laser power during 2 s in a 6 pixels radius circle. Data analysis was performed with the ImageJ software by measuring the displacement (distance and orientation) reached at the maximal recoil of all vertices in a 10 μm radius circle around the ablated vertex, from their initial location (before cut), using Manual tracking plugin.

For circular photo-ablation, the focal plane was illuminated during 4-5 s along a 45 pixels radius circle (Fig7f-i). This focal plane was acquired every 0.551 s, during 10 s before and at least 45 s after ablation, with a pixel size of 0.13 μm/pixel. For the second set of experiments (wound healing), tissue was first wounded at the level of a tricellular junction by illuminating during 4-5 s along a 10 pixels radius circle and then cut circularly during the healing phase using the circular shape described above (Fig7f-i). Data analysis was performed with the ImageJ software. Briefly, the distance between the location of the circular cut and the maximal recoil induced by this ablation was measured at 0° (proximal-distal axis) and 90° (circular axis) using the line tool.

### The Vertex Model

#### Initial tissue generation

We modeled the most distal part of the leg imaginal disc (Fig1a), before the T4-T5 fold formation begins, as a 2D meshwork around a cylinder capped by two hemispheres, oriented along the proximal-distal axis. We started with a mostly hexagonal lattice with a perimeter of 23 cells and a length of 15 cells. We perform two rounds of cell divisions with a random division axis to randomize cell side number and create a meshwork of approximately 50 by 30 cells. Diversity of cell shapes is increased by adding variability to the cells’ preferred areas, normally distributed with an 8% variance. The initial tissue has 1652 cells, is 200 μm long in its proximal-distal axis and 100 μm in diameter.

#### Mechanical model

The epithelium shape is given by the quasi-static equilibrium of a potential energy dependent on the junctional mesh geometry, following our previous work (Gracia et al., 2019; Monier et al., 2015).

This energy is given in equation (1) and is comprised of three cell-level terms and two constrain terms. At the cell level, apical shape is governed by area and perimeter elasticity terms, following (Bi et al., 2015). To these terms, we add a linear apico-basal tension for the apoptotic cells, dependent on cell height. Two more terms ensure maintenance of the overall tissue shape. First, the total volume of the tissue is maintained by an elastic constrain. Second, an external barrier is modeled as a sphere surrounding the tissue; when the distance of a vertex from the sphere center is higher than the sphere radius, it is pulled back to this radius by an elastic force.

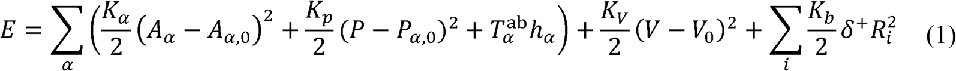

The value of (KV) was chosen at the lowest value such that compression of the tissue by cellular contractility and the effect of the lumen on the fold formation was minimal and kept the tissue integrity (see FigS6). Here ∑_α_ marks the sum over every polygonal cell and ∑_*i*_ a sum over every vertex. Apico-basal tension is exerted from the vertices towards the proximal-distal axis (center of the cylinder). We consider an anchor point *i*′ as the protection of the vertex *i* onto the proximal-distal axis. *i*′ is rigidly fixed to this axis. The penetration depth *δ*^+^ *R_i_* is defined by:

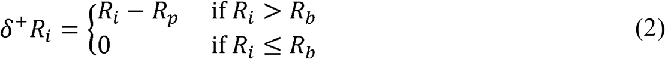

Energy minimum is reached through a gradient descent strategy using the Broyden-Fletcher-Goldfarb-Shanno bound constrained minimization algorithm from the scipy library (van der Walt et al., 2011).

#### Tissue anisotropy

In vivo, cell polarity translates in different mechanical properties for different cell junctions. Here, in order to create the cell anisotropy (the ratio between the long axis and the perpendicular axis of the apical surface of the cell), we added a weight on each cell junction in the calculation of cell perimeter. The modified perimeter is calculated as the weighted sum

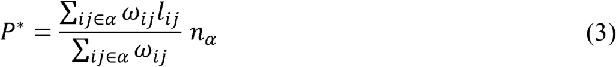

Where ∑_ij∈α_ denotes a sum over all edges of cell *α*, and *ω_ij_* is the weight of the junction *ij*, *l*_*ij*_ is the length of the junction *ij* and *n_α_* is the number of sides of cell *α*. Note that this weighted sum is equal to the actual perimeter when all weights are equals, and allows to model polarity without any modification of the other dynamical parameters.

We set two different values of the weight depending on the orientation of the edge. Weight is higher for circumferential edges (*ω*_//_) than for proximal-distal ones (*ω*_⊥_). (FigS7c-h). As cell shapes are on average hexagonal, we set the boundary between parallel and perpendicular junctions at *π*/3.

#### Apoptotic process

Around 30 apoptotic cells are chosen randomly in the fold region according to a probability density described in our previous work (Monier et al., 2015). The tissue deforms progressively as apoptotic cells undergo apical constriction and apico-basal traction through gradual changes in their mechanical parameters, while other cells passively follow the deformation.

A cell starts apoptosis at time step *t_i_* such that 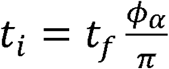 where *ϕ_α_* is the angle of the cell’s apical surface face center with respect to the dorso-ventral axis and *t_f_* the final simulation time.

Apoptosis is modeled as a sequence of apical constriction and apico-basal traction.

Apical constriction consists in a reduction of the cell’s preferred perimeter *P*_*α*,0_ at a constant rate 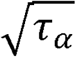 and a reduction of its preferred area *A*_*α*,0_ at constant rate *τ*_α_ until the cell area reaches a threshold *A_c_* as given in equation (4). Once the critical area is reached, preferred area and preferred perimeters are maintained constant.

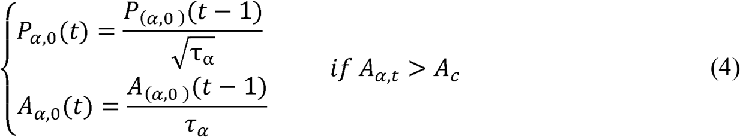

During the apical constriction phase, contraction is propagated to neighboring cells. Contraction rate *τ_β_* of a neighboring cell *β* decreases linearly as the cell is farther away from the apoptotic cell:

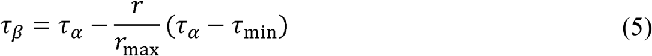

Here *r* = 1 if the cell *β* is a direct neighbour of the apoptotic cell, 2 if it’s a second order neighbor, and so on. *r*_max_ is the span of the propagation and *τ_min_* the contraction rate for cells at *r_max_* from the apoptic cell.

Each apoptotic cell can develop an apico-basal tension, during and after the constriction phase, with probability *p* = exp(−*A*_α_/*A*_c_). Traction takes place for *N*_t_ time steps, during which the apico-basal tension exerted on the face is *T*^ab^

#### Mechanical perturbation

We modeled mechanical perturbations during fold formation as three “disrupting cells” placed randomly at a maximal distance of 40 μm (7 cells distance) from the middle of the predicted fold position. The “disrupting cell” dynamics is similar to apoptosis, with apical constriction and apical-basal tension, however, apical constriction starts at the beginning of the simulation, and when the critical area is reached, apico-basal traction is applied until the end of the simulation. For every simulation, apart from the time span for which the mechanical perturbation is exerted, the parameters for the perturbing cell are identical to the parameters for an apoptotic cell.

#### Choice of parameter values

The unit energy (denoted by *u*) is defined so that the area elasticity modulus *K_α_* equals 1*u*/*μm*^4^ To model lumen incompressibility, lumen volume elasticity *K_V_* is such that apical contraction compresses the super-ellipsoid by 5% in volume (*K_V_* = 1.10^−5^*u*/μm^6^ (FigS4). Preferred area *A*_0_ and preferred perimeter *P_0_* are chosen to have a constant 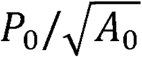 ratio of 2 throughout the simulation, this corresponds to a stiff tissue in (Bi et al., 2015) framework. With the above value of 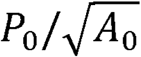, we choose the apical perimeter elasticity *K*_p_ to allow cell shape changes upon apical constriction.

The code used for modelling is publicly available: https://github.com/DamCB/tyssue and https://github.com/suzannelab/polarity.

### Quantification and statistical analysis

#### Quantitative analysis of the fold variability

We characterized the variability of fold formation *in vivo* by measuring the maximal distance between the real fold from the predicted fold, the angle of the fold relative to the proximal-distal axis and the parallelism between folds, for *control* and *Arpc5 RNAi* expressing leg discs (genotypes are indicated in each figure legends), as follow:

The maximal distance between real fold and predicted fold was measured in two steps. First, we measured the distance between the Notch signaling (labeled by Deadpan immunostaining) and the position of maximal deviation of the fold using the straight-line tool in ImageJ. Second, we calculated the mean of maximal distance in the control and subtracted the value obtained to each measure to normalize them independently from the genotype. Standard deviation of control legs was measured and used to define predicted fold (± 1.30 μm from the mean), which is highlighted in yellow on the graph in Figure 1c. Because the width of the *apterous* domain is more variable in AJs GFP-trap experiments, in Figure 6d we measured the relative maximal distance rather than the absolute maximal distance, by dividing the distance between the T3-T4 and the T4-T5 fold by the width of the *apterous* domain.

- The angle formed between the fold and the proximal-distal axis, was measured in ImageJ using the angle tool.
- The fold parallelism was calculated as the ratio between angles of two folds of the same leg.

#### Quantitative analysis of Dpn and Rpr expression domain

Dpn and Rpr stripes of expression have been characterized using Image J software. The stripes have been outlined to define ROIs. From each ROI, the main axis (or ferret) has been defined and the orientation of this axis relative to the proximo-distal axis defined. Then, the voronoi (corresponding to the points equidistant from the proximal and distal border of the domain) was obtained to determine the mean width of the domain and the curvature.

#### Cartography of mechanical perturbations *in vivo*

Zones of high apico-basal tension, or “mechanical perturbations”, were spotted on 3D reconstructions of (*sqh-GFP[29B]; Dll-Gal4*) and (*sqh-GFP[29B]; Dll-Gal4; UAS-Arpc5RNAi*) leg discs using Imaris and located on corresponding positions on rolled-out maps of the fold regions (Fig3f). To each perturbation corresponds a set of coordinates defined (in x) by the angle formed between the line going from the center of the leg to the perturbation and the DV axis and (in y) by the distance between the perturbation and the predicted fold.

#### Analysis of Junctional Intensities

Using the surface tool on Imaris (Bitplane), a mask was created from the junctional labelling (arm-GFP, α-catenin-RFP or E-cadherin antibody) to quantify Myosin II, F-actin, adherens junctions or Arp3 present at junctions. From this new Z-stack, the maximum of intensity (MaxProj) and the sum of intensity (SumProj) were projected using Fiji (ImageJ 1.51s – NIH). Then, from the MaxProj, a skeleton was created and each junction was individualized by suppressing nodes. This template was used to identify each junction as a region of interest (ROI). From these ROIs, the angle – formed with the proximal-distal axis – and the raw fluorescence intensity of each junction were measured on the SumProj (allowing to sum the full quantity of Myosin II present at a given xy position). Manual correction was done if required.

The junctional mean of fluorescence intensity 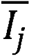 along junction was normalized to the average junction intensity over all the junctions analyzed (fold+adjacent domain) 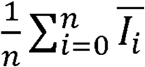

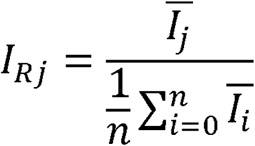

#### Analysis of the cell aspect ratio

Using Fiji software, the maximal intensity projection was done from a Z-stack and a skeleton was created. This template was used to identify each cell as a region of interest (ROI). From these ROIs, the aspect ratio (major axis / minor axis) was measured in order to define the anisotropy of the cell.

#### Quantitative analysis of the apical myosin

Using ImageJ software, a segmented line was drawn along the apical surface of the posterior (hh-Gal4; UAS-GFP) and the anterior (control) domains on five different z sections for each leg. For each z section, the mean of fluorescence intensity was measured and the ratio between posterior and anterior domains was calculated. The mean of these five ratios per leg is represented as one spot on the graph in Figure S5a.

#### Quantitative analysis of medio-apical myosin flow

Particle image velocimetry from time-lapse movies of E-cadh-GFP; sqh-RFPt leg discs was quantified using the OpenPIV Python package. Images were acquired on a Zeiss LSM 880 using High Resolution Airy Scan with a time frame of 15”. Flows were analyzed in 2D on maximum projection of standard deviation. Each cell was isolated using Image J software. The displacement of particles was tracked between successive frames and the mean displacement is presented in a polar charts in FigS5e-f. n=14 for control and n=20 for Arpc5 RNAi.

#### Cartography of mechanical perturbations *in silico*

In the simulations, each cell has assigned coordinates, allowing us to spot the positions of each mechanical perturbation (randomly generated in the model) in a rolled-out map of the fold domain.

#### Measure of *in silico* fold deviation

We characterized the variability of fold orientation in the different sets of simulations: with or without mechanical perturbations and with or without tissue polarity. For each simulation, we spotted the position of maximal depth automatically as the point located at the minimal distance from the central axis of the tissue. We then defined its coordinates in relation to the predicted fold position and the DV axis. To calculate the real maximal deviation from the predicted fold domain, we subtracted the maximal deviation obtained without perturbation from the one obtained with perturbations for a given pattern of apoptosis. To calculate the angle formed between the fold and the proximal-distal axis, we measured the angle between the predicted fold and the line formed from fold position on the ventral side and the position of maximal depth. To calculate the real maximal angle from the predicted fold domain, we subtracted the angle without perturbation from the one obtained with perturbations for a given pattern of apoptosis.

#### Statistical Analysis

The normality of the data sets was determined using Prism 8 (Graph Pad).

A Mann-Whitney U-test was used to assess the significance of differences in tissue recoil after laser ablation (Fig2i; Fig3c,e; Fig7h-i and FigS7a-b) or in apical intensity of MyoII (FigS5a), considering legs as independent from each other. The null hypothesis was that measurements were samples from the same distribution. Tests were performed using Prism 8 (Graph Pad).

Variances of the phenotypes observed *in vivo* were compared using the F-test of equality of variances in Prism 8 (Graph Pad), considering that different data sets follow a normal distribution. Variances from simulated data were compared using the Levene’s test in Python 3.7, considering that different data sets do not fit with a normal distribution. The null hypothesis was that variances of population were equal.

Spearman correlation coefficients (FigS7g-h) and associated p-values were computed online (www.wessa.net/rwasp_spearman.wasp).

##### Supplemental video titles

**Movie 1.** 3D reconstruction of a control leg disc showing the expression of Deadpan and the fold domain (related to Fig2).

**Movie 2.** 3D reconstruction of an Arpc5 RNAi leg disc showing the expression of Deadpan and the fold domain (related to Fig2).

**Movie 3.** Laser ablation experiments of apoptotic myosin II cables in control and Arpc5 RNAi leg discs (Related to Fig2).

**Movie 4.** Laser ablation experiments of non-apoptotic apico-basal structures of Myosin II in control and Arpc5 RNAi leg discs (Related to Fig3).

---

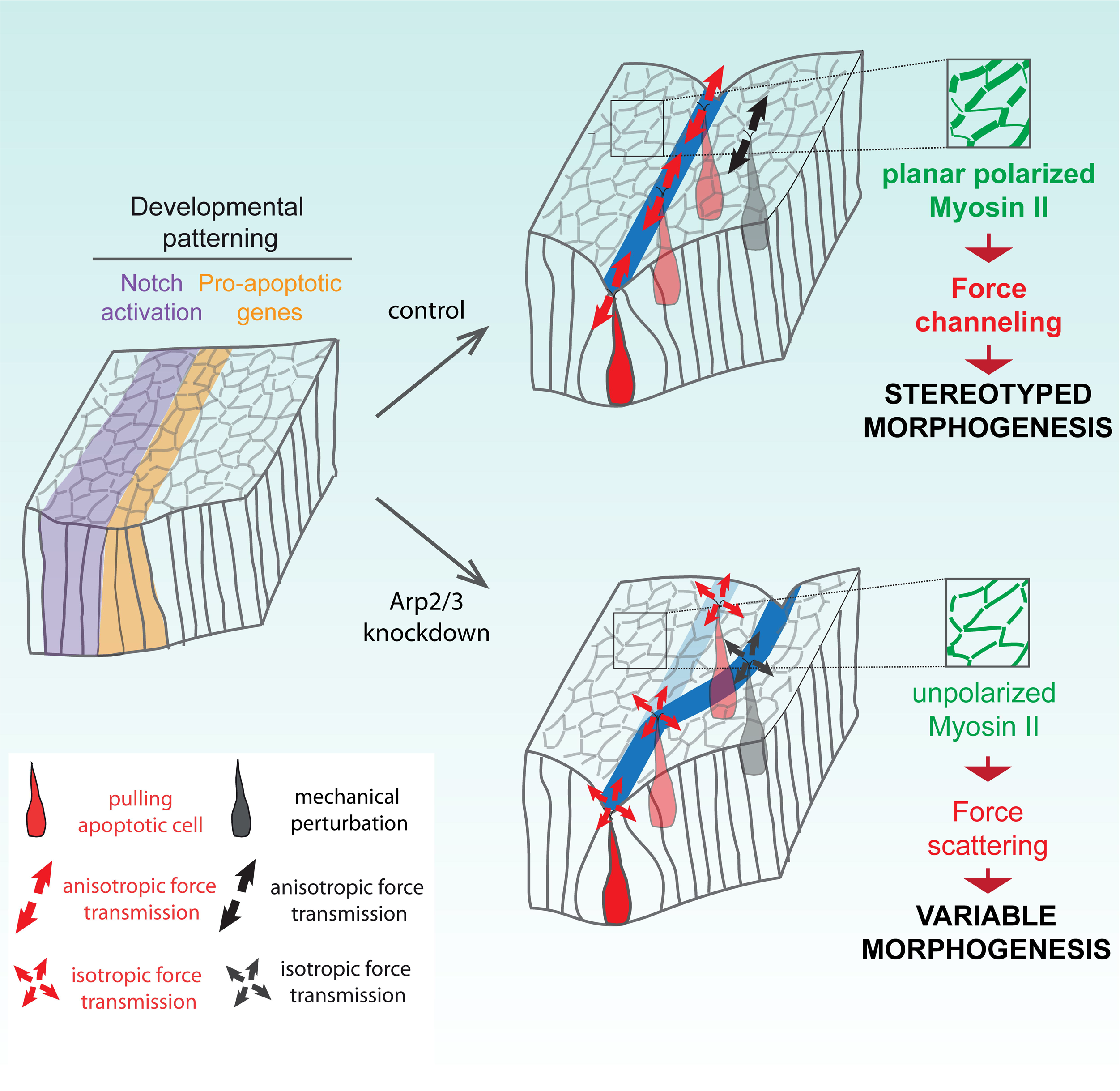

## Notes

### Competing Interest Statement

The authors have declared no competing interest.

