## Supplementary material for "Mechanical control of morphogenetic robustness in an inherently challenging environment": Sup Fig

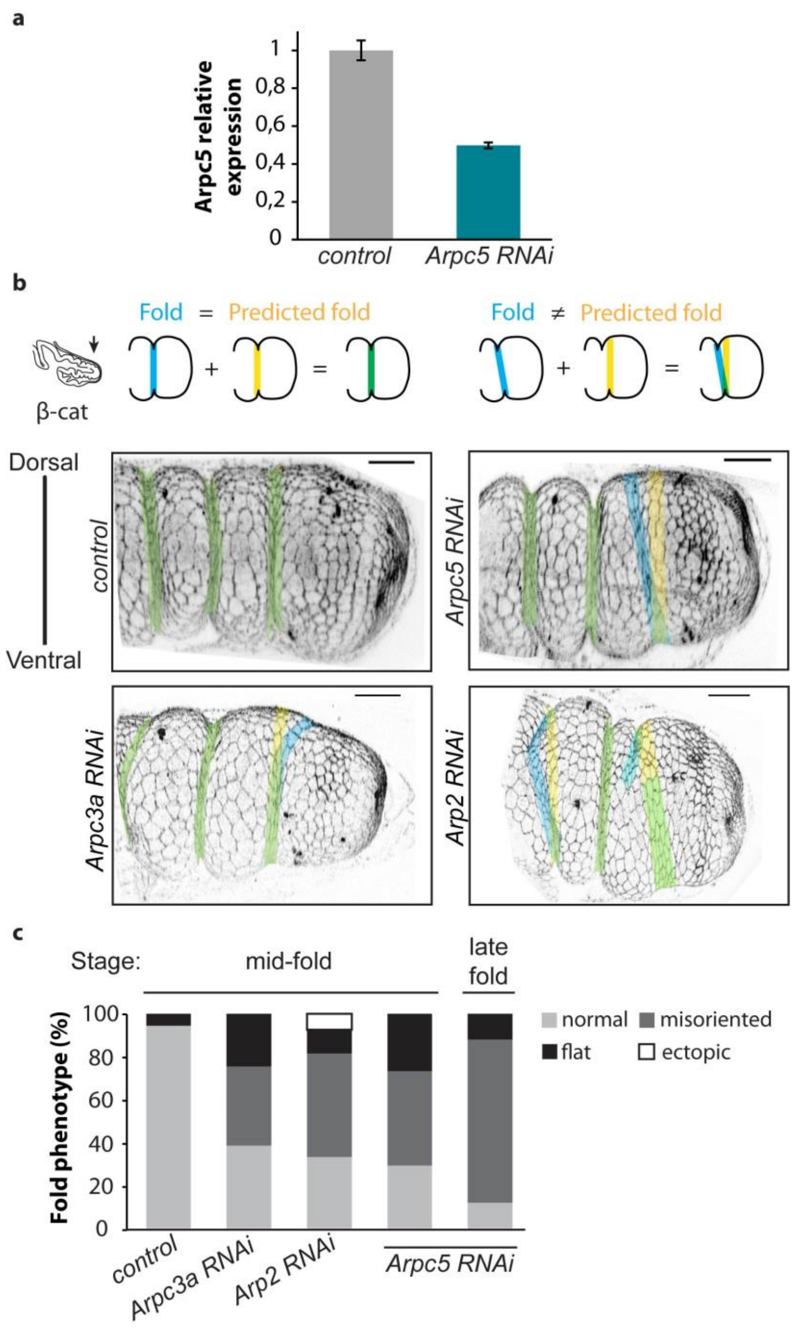


**Figure S1: Fold defects in Arp2/3 knockdown conditions (Related to Figure 1).**

**a,** Normalized expression of Arpc5 mRNA in *control* and *Arpc5 RNAi*. Bar graphs indicate the mean ± SEM.

**b,** 3D reconstructions of β-catenin-GFP (*arm-GFP*) leg discs (bottom) and corresponding schematics (top) showing fold phenotypes in *control*, *Arpc5* RNAi, *Arpc3a* RNAi and *Arp2* RNAi conditions. Predicted folds are highlighted in yellow, real folds in blue and perfect match between them in green. Scale bar represents 20 µm.

**c,** Quantification of T4-T5 fold phenotypes in *control*, *Arpc3a* RNAi *Arp2* RNAi and *Arpc5* RNAi leg discs at mid-fold (WP+1h30 at 29°C) and late fold stage (WP+2h40 at 29°C) (n=54, 57, 27, 60, 25 legs, respectively).

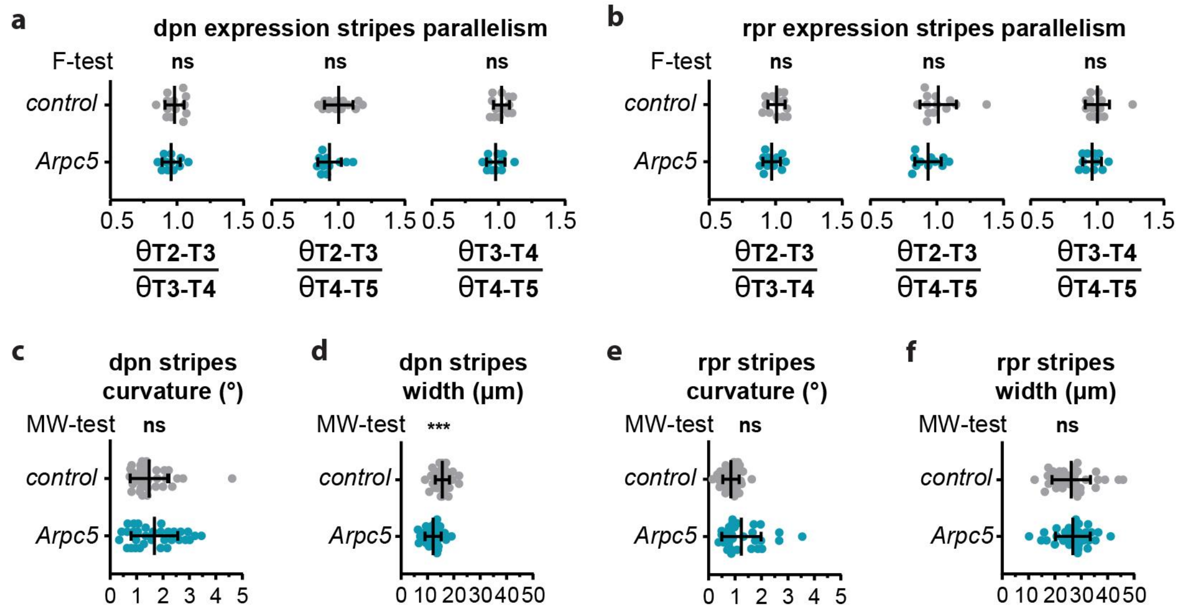


**Figure S2: (related to Figure 2).**

**a-b**, Dot plots quantifying the parallelism of the stripes of expression of deadpan (e) or reaper (b) in *control* (*rpr-lacZ; Dll-Gal4*) and *Arpc5* RNAi (*rpr-lacZ; Dll-Gal4; UAS-Arpc5RNAi*) (n=13 and 11 legs respectively) leg discs. A *F*-test of equality of variances has been used to compare the phenotypic variances between *control* and *Arpc5 RNAi* pupae. ns, not significant. Black lines represent the mean ± SD.

**c-f**, Dot plots showing the curvature (c,e) or the width (d,f) of the stripes of expression of deadpan (c,d) or reaper (e,f) in *control* (*rpr-lacZ; Dll-Gal4*) and *Arpc5* RNAi (*rpr-lacZ; Dll-Gal4; UAS-Arpc5RNAi*) (n=13 and 11 legs respectively) leg discs. Black lines represent the mean ± SD**.** Statistical significance has been calculated using Mann Whitney U test. ns, not significant; ***, p-value < 0.001.

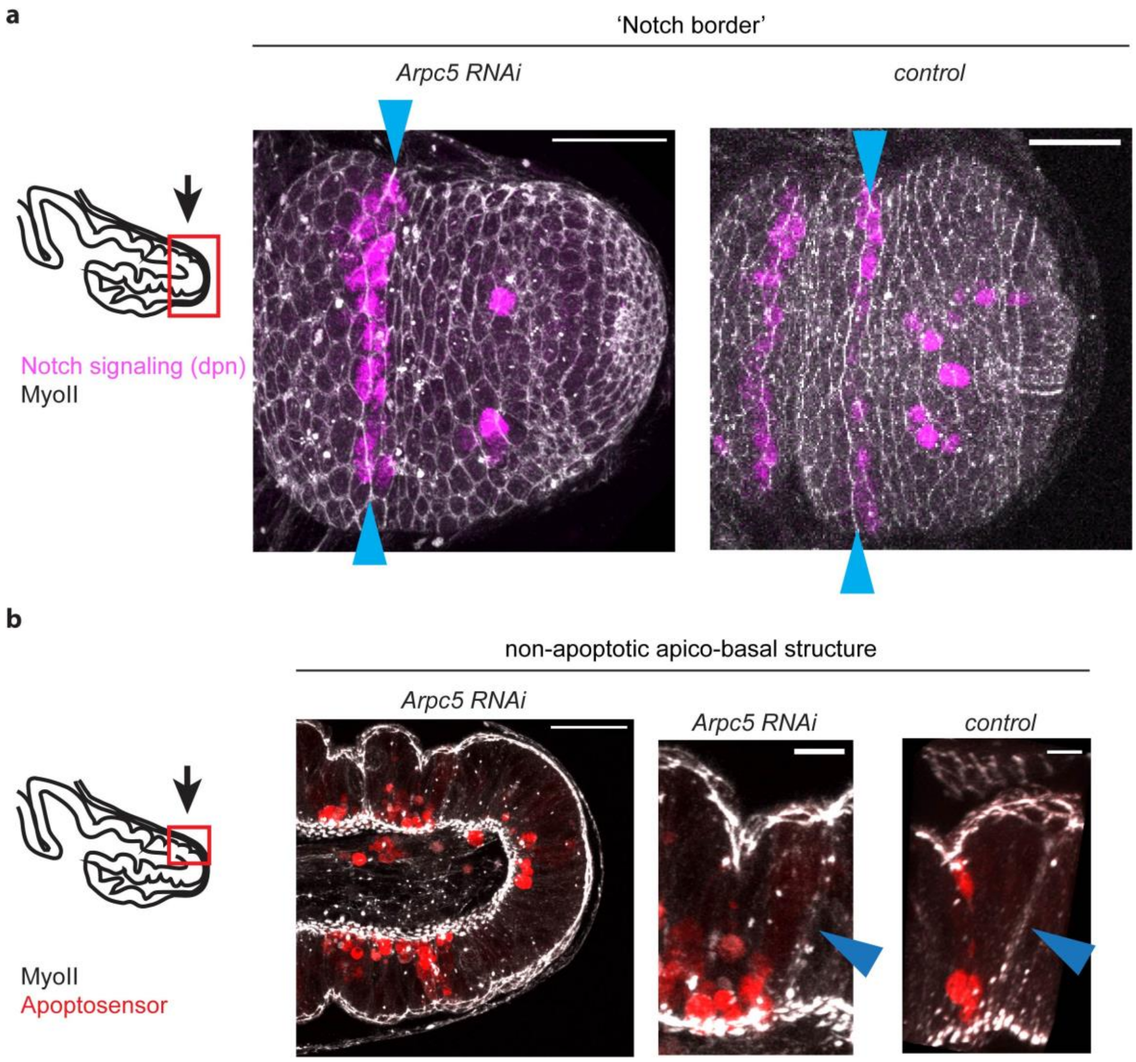


**Figure S3: Characterization of mechanical perturbations (Related to Figure 3).**

**a,** Confocal images of *Arpc5* RNAi (*sqh-GFP[29B]; Dll-Gal4; UAS-Arpc5RNAi*) and *control* (*sqh-GFP[29B]; Dll-Gal4*) leg discs stained with an anti-deadpan antibody (magenta) showing that the supra-cellular cable of myosin is formed at the border of Notch activation domain. Blue arrowheads indicate the Notch activation domain. Scale bar represents 20 µm.

**b,** Sagittal views of *Arpc5* RNAi (*sqh-RFPt[3B]; Dll-Gal4, UAS-GC3Ai; UAS-Arpc5RNAi*) and *control* (*sqh-RFPt[3B]; Dll-Gal4, UAS-GC3Ai*) leg discs showing that the apico-basal structures of Myosin II observed outside the fold domain (blue arrowheads) are not related to the presence of apoptotic cells (red). Scale bar represents 20 µm and 5 µm, respectively for general and close-up view.

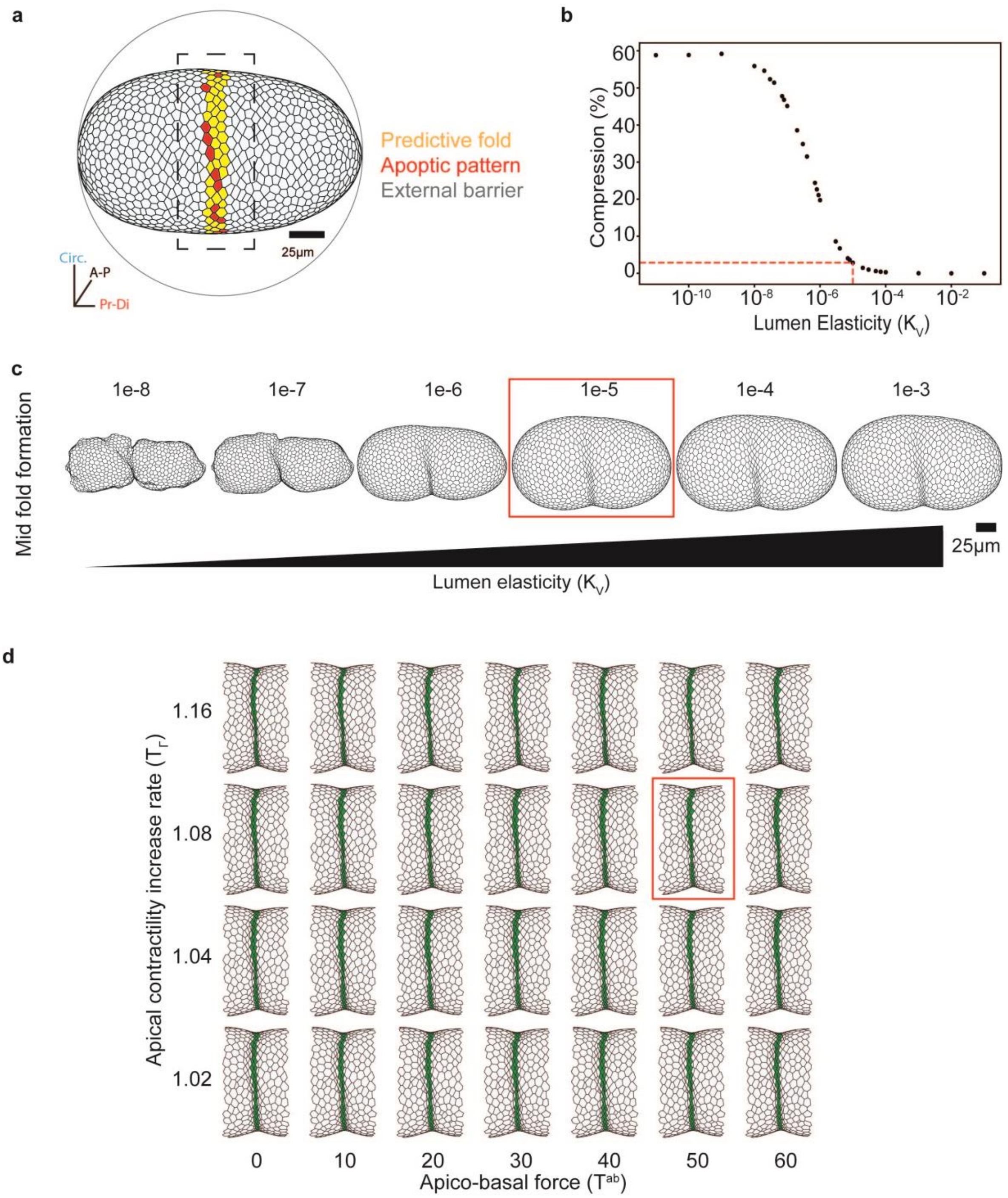


**Figure S4: Simulations of epithelium folding (Related to Figure 3).**

**a,** Lateral view of the virtual tissue. Cells from the predicted fold domain are indicated in yellow, apoptotic cells in red. The tissue is constrained by a spherical external barrier represented by the grey circle. The black dotted square corresponds to the close-up shown in **d**.

**b,** Relative change in the virtual tissue volume as a function of lumen elasticity $K^V$. Compression is expressed as the ratio between the observed volume and the equilibrium volume V_L^0.

**c**, Aspect of the virtual tissue at maximum fold depth for different values of $K^V$. Red squared corresponds to the choosen value for $K^V$.

**d,** Simulations obtained at maximal fold depth for distinct values of apical contractility increase rate (y-axis) and apico-basal force (x-axis). All other parameters are unchanged. Fold is in green.

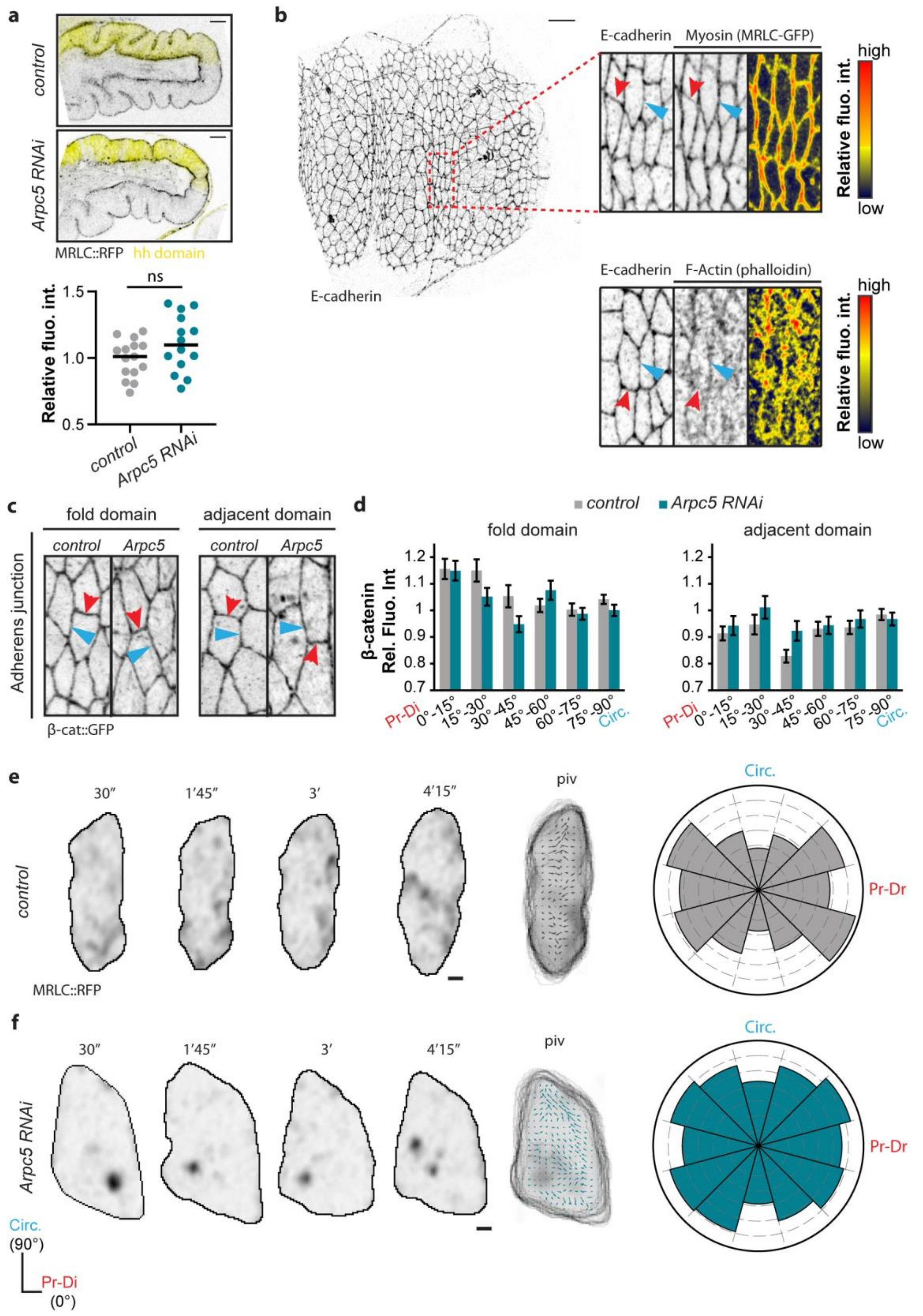

a
control
Arpc5 RNAi
MRLC::RFP hh domain
ns
Relative fluo. int.
1.5
1.0
0.5
control
Arpc5 RNAi
b
E-cadherin
E-cadherin Myosin (MRLC-GFP)
high
Relative fluo. int.
low
E-cadherin F-Actin (phalloidin)
high
Relative fluo. int.
low
c
fold domain
adjacent domain
control Arpc5
control Arpc5
Adherens junction
β-cat::GFP
d
control Arpc5 RNAi
fold domain
adjacent domain
β-catenin Rel. Fluo. Int
1.2
1.1
1
0.9
0.8
0.7
Pr-Di 0°-15° 15°-30° 30°-45° 45°-60° 60°-75° 75°-90° Circ.
e
30″ 1′45″ 3′ 4′15″
piv
control
MRLC::RFP
Circ.
Pr-Dr
f
30″ 1′45″ 3′ 4′15″
piv
Arpc5 RNAi
Circ.
Pr-Dr
Circ. (90°)
Pr-Di (0°)

**Figure S5: Cortical Myosin II intensity level and adherens junctions in Arpc5 knockdown (Related to Figure 4)**

**a**, (top) Confocal images showing Myosin II distribution in a control (*sqh-GFP; hh-Gal4, UAS-LacZ*) (top, n=15) or a *Arpc5* RNAi (*sqh-GFP; hh-Gal4, UAS-Arpc5RNAi*) (bottom ; n=14) leg discs. The posterior "Hedgehog" domain is highlighted in yellow. Scale bar represents 30 µm. (bottom) Dot plots showing the relative fluorescence intensity of Myosin II calculated as the ratio between the posterior domain (hh domain, in yellow) and the anterior domain.

**b,** (left) Confocal images showing the leg morphogenesis in *sqh-RFPt; Dll-Gal4; UAS-Arp3-GFP* animals. Scale bar represents 10 µm. (right) Close up views of confocal images showing the distribution of adherens junction (E-cadherin) and either Myosin II (top) or F-actin (bottom, stained with phalloidin). The intensities of myosin and actin are color-coded as indicated. Red and blue arrowheads indicate proximal-distal and circumferential junctions, respectively

**c**, Close up views of confocal images showing the distribution of adherens junctions in control (Dll-Gal4; arm-GFP) or Arpc5 RNAi (Dll-Gal4; arm-GFP, UAS Arpc5RNAi) leg discs in the fold domain (left) and in the adjacent domain (right). Red and blue arrowheads indicate proximal-distal and circumferential junctions, respectively

**d**, Quantification of β-catenin distribution at junctions in leg discs controlor Arpc5 RNAi, in both the fold domain (left; n= 668 and n= 571 junctions respectively) and the adjacent domain (right; n= 578 and n= 449 junctions respectively). Bar graphs indicate the mean ± SEM.

**e-f**, PIV analysis of MRLC-RFP flows in control (**d**) and Arpc5 RNAi (**e**). Stills from isolated cells are presented on the left showing Myosin II dynamics, with vectors representing the local displacements superimposed on a time projection. On the right, polar charts represent the mean distance of local displacements for each orientation.

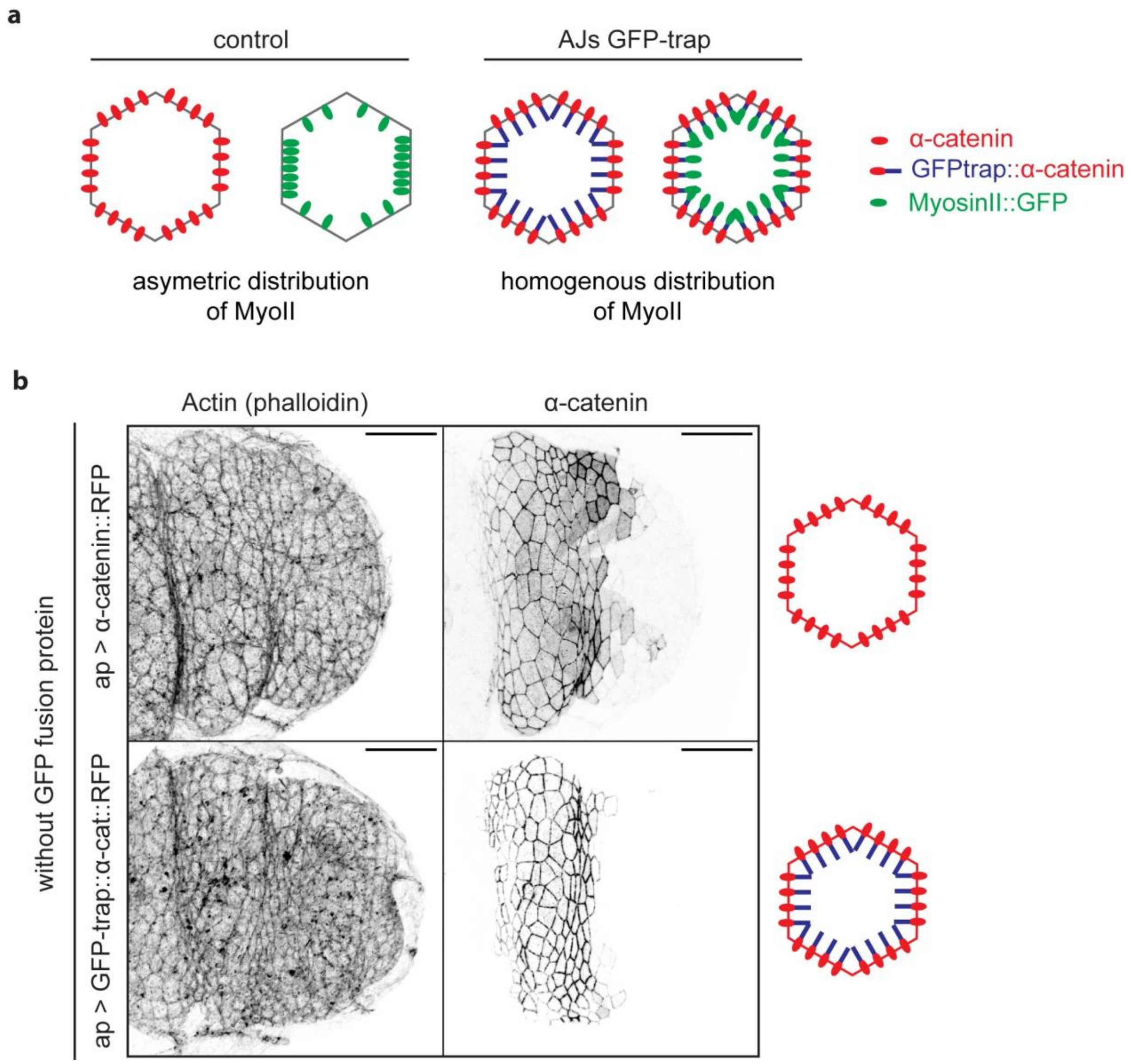


**Figure S6: Characterization of the AJs-GFP-trap construct (Related to Figure 6).**

**a,** Schematics of junctional Myosin II distribution in a *control* or an AJs-GFP-trap; MyosinII-GFP conditions.

**b,** Confocal images of *tubG80ts*; *ap-Gal4, UAS-α-catenin-TagRFP* and *tubG80ts; ap-Gal4, UAS-vhhGFP4-α-catenin-TagRFP (AJs-GFP-Trap)* leg discs in the absence of GFP fusion protein. The F-actin distribution (phalloidin) shows that the expression of AJs-GFP-Trap alone does not perturb epithelial organization. Scale bar represents 20 µm.

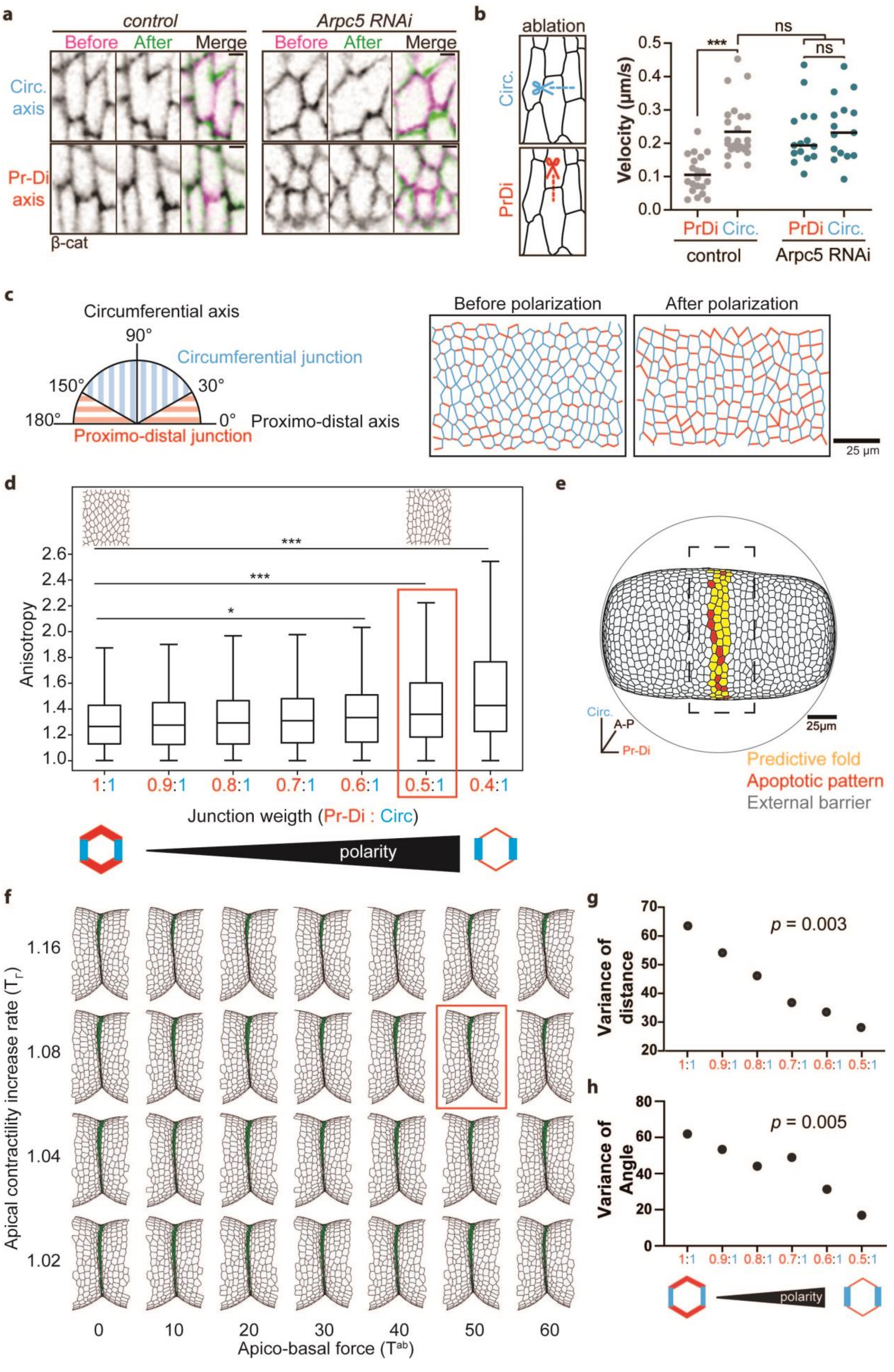

a
control
Arpc5 RNAi
Before After Merge
Circ. axis
Pr-Di axis
β-cat
b
ablation
Circ.
PrDi
Velocity (μm/s)
***
ns
PrDi Circ. PrDi Circ.
control
Arpc5 RNAi
c
Circumferential axis
90°
Circumferential junction
150°
30°
180°
0°
Proximo-distal axis
Proximo-distal junction
Before polarization
After polarization
25 μm
d
Anisotropy
1:1 0.9:1 0.8:1 0.7:1 0.6:1 0.5:1 0.4:1
Junction weigth (Pr-Di : Circ)
polarity
e
Circ.
A-P
Pr-Di
25μm
Predictive fold
Apoptotic pattern
External barrier
f
Apical contractility increase rate (Tr)
1.16
1.08
1.04
1.02
0 10 20 30 40 50 60
Apico-basal force (Tab)
g
Variance of distance
p = 0.003
h
Variance of Angle
p = 0.005
polarity

**Figure S7: Simulations of fold formation with different values of planar polarity of junctional tension. (Related to Figure 6).**

**a,** Confocal images showing the dynamics of adherens junctions before and after laser ablation of circular (top) and proximal-distal (bottom) junctions in *control* (*Dll-Gal4; arm-GFP*) and *Arpc5 RNAi* (*Dll-Gal4; arm-GFP, UAS-Arpc5RNAi*) leg discs. Scale bar represents 2 µm.

**b,** Dot plots showing initial recoil velocity of junctions after laser microdissection in *control* and *Arpc5* RNAi leg discs (n=24, 20, 15 and 15 respectively). Statistical significance has been calculated using Mann Whitney U test. ns, not significant; ***, p-value < 0.001.

**c,** Left panel: Scheme representing the distribution of circumferential and proximal-distal junctions according to their orientation. Blue junctions are considered oriented in the circumferential axis. Red junctions are considered oriented in the proximal-distal axis. Right panel: repartition of the two types of junctions in the simulation with or without polarization of the tissue.

**d,** Box plots presenting the values of tissue anisotropy for different junction weight ratios (weight of proximal-distal junctions/weight of circumferential junctions). The preferred junctional weight ratio is indicated by the red square. Cell shapes are no longer physiological for lower values of weight ratio (data not shown).

**e,** Simulation showing the general organization of the polarized tissue before fold formation. Cells from the predicted fold are in yellow, apoptotic cells in red. The tissue is surrounded by a spherical external barrier represented by the grey circle. The black dotted square corresponds to the close-up in panel **g**.

**f,** Simulations obtained at maximal fold depth for distinct values of contractility increase rate (y-axis) and apico-basal force (x-axis) with planar polarity. All other parameters are unchanged. Fold is in green. Preferred values are indicated by a red square.

**g,** Scatter plot showing the correlation between the variance of the fold deviation and the polarity of the tissue from 55 in silico analysis. A Spearman rank correlation has been used to evaluate this correlation. The p-value is indicated under the graph.

**h,** Scatter plot showing the correlation between the variance of the fold angle and the polarity of the tissue from 55 in silico analysis. A Spearman rank correlation has been used to evaluate this correlation. The p-value is indicated under the graph.
